# Large-scale biophysically detailed model of somatosensory thalamocortical circuits in NetPyNE

**DOI:** 10.1101/2022.02.03.479029

**Authors:** Fernando S. Borges, Joao V.S. Moreira, Lavinia M. Takarabe, William W. Lytton, Salvador Dura-Bernal

## Abstract

The primary somatosensory cortex (S1) of mammals is critically important in the perception of touch and related sensorimotor behaviors. In 2015, the Blue Brain Project developed a groundbreaking rat S1 microcircuit simulation with over 31,000 neurons with 207 morpho-electrical neuron types, and 37 million synapses, incorporating anatomical and physiological information from a wide range of experimental studies. We have implemented this highly-detailed and complex S1 model in NetPyNE, using the data available in the Neocortical Microcircuit Collaboration Portal. NetPyNE provides a Python high-level interface to NEURON and allows defining complicated multiscale models using an intuitive declarative standardized language. It also facilitates running parallel simulations, automates the optimization and exploration of parameters using supercomputers, and provides a wide range of built-in analysis functions. This will make the S1 model more accessible and simpler to scale, modify and extend in order to explore research questions or interconnect to other existing models. Despite some implementation differences, the NetPyNE model preserved the original cell morphologies, electrophysiological responses and spatial distribution for all 207 cell types; and the connectivity properties of all 1941 pathways, including synaptic dynamics and short-term plasticity (STP). The NetPyNE S1 simulations produced reasonable physiological firing rates and activity patterns across all populations. The network generated a 1 Hz oscillation comparable to the original model in vitro-like state. By then reducing the extracellular calcium concentration, the model reproduced the original S1 in vivo-like states with asynchronous activity. These results validate the original study using a new modeling tool. Simulated local field potentials (LFPs) exhibited realistic oscillatory patterns and features, including distance- and frequency-dependent attenuation. The model was extended by adding thalamic circuits, including 6 distinct thalamic populations with intrathalamic, thalamocortical and corticothalamic connectivity derived from experimental data. The thalamic model reproduced single known cell and circuit-level dynamics, including burst and tonic firing modes and oscillatory patterns, providing a more realistic input to cortex and enabling study of thalamocortical interactions. Overall, our work provides a widely accessible, data-driven and biophysically-detailed model of the somatosensory thalamocortical circuits that can be employed as a community tool for researchers to study neural dynamics, function and disease.

## 1 Introduction

The primary somatosensory cortex (S1) of mammals is critically important in the perception of touch and works closely with other sensory and motor cortical regions in permitting coordinated activity with tasks involving grasp (Petrof, Viaene, and Sherman 2015; Barthas and Kwan 2017; Bosman et al. 2011). Moreover, the communication of these cortical areas with the thalamus is crucial for maintaining functions, such as sleep and wakefulness, considering that the thalamocortical (TC) circuit is essential for cerebral rhythmic activity (O’Reilly et al. 2021). A greater understanding of S1 cortical circuits will help us gain insights into neural coding and be of assistance in determining how disease states such as schizophrenia, epilepsy and Parkinson’s disease lead to sensory deficits or uncoordinated movement (Petrof, Viaene, and Sherman 2015; Vázquez, Salinas, and Romo 2013; Azarfar et al. 2018; Peña-Rangel et al. 2021).

There exists an impressive, highly-detailed model of rat S1 developed by the Blue Brain Project (BBP) (Markram et al. 2015), incorporating anatomical and physiological information from a wide range of experimental studies. This groundbreaking model includes over 31,000 neurons of 55 layer-specific morphological and 207 morpho-electrical neuron subtypes, and 37 million synapses capturing layer- and cell type-specific connectivity patterns and synaptic dynamics. Simulation results matched in vitro and in vivo experimental findings, and the model has been used over the years to reproduce additional experimental results and generate predictions of the dynamics and function of cortical microcircuits (Michael W. Reimann et al. 2015; Gal et al. 2017; Michael W. Reimann, Horlemann, et al. 2017; Michael W. Reimann, Nolte, et al. 2017; Amsalem et al. 2020; Hagen et al. 2018). Although the BBP S1 model is state-of-the-art, certain constraints limit its reproducibility and use by the community, as well as its extension or modification to connect to other regions or update model features. The size and complexity of any model of this scope is daunting. Due to its scale and complexity, the original model must be run and analyzed on large High Performance Computing platforms (HPCs), which are not available to many users. Although the model is simulated using NEURON (Carnevale and Hines 2006; Migliore et al. 2006) a widely used platform within the computational neuroscience community, it also requires other custom libraries specifically designed to facilitate this workflow. These libraries are used to build, manage simulations and analyze the model. However, not all of these libraries and workflows are publicly available (Markram et al. 2015), making it somewhat difficult to modify the code, and scale or simplify the model for simulation on smaller computers, overall reducing its accessibility and reproducibility (McDougal, Bulanova, and Lytton 2016).

Here we implemented the original BBP S1 model in NetPyNE (Dura-Bernal et al. 2019) in order to make it more accessible and simpler to scale, modify and extend. NetPyNE is a python package that provides a high-level interface to the NEURON simulator, and allows the definition of complex multiscale models using an intuitive declarative standardized language. NetPyNE translates these specifications into a NEURON model, facilitates running parallel simulations, automates the optimization and exploration of parameters using supercomputers, and provides a wide range of built-in analysis functions.

Conversion to NetPyNE also makes it easier to connect to previous models developed within the platform, such as our primary motor cortex model (Sivagnanam et al. 2020; Dura-Bernal, Neymotin, et al. 2022), and models implemented in other tools (e.g. NEST) by exporting to the NeuroML or SONATA standard formats. In prior work, we ported a classic model of generic sensory cortical circuits (Potjans and Diesmann 2014) to our NetPyNE platform (Romaro et al. 2021) in order to make it both more scalable and facilitate modification of cell models and network parameters. The original model used integrate-and-fire neurons and we replaced these with more complex multi-compartment neuron models.

Although we have primarily focused on simplifying the network description, we have also made the model more complex, and more complete, by adding the associated somatosensory thalamic circuits and bidirectional connectivity with cortex to allow interplay of these two highly coordinated areas (Meyer et al. 2010). The deepening of knowledge about the cortico-thalamo-cortical loop (Shepherd and Yamawaki 2021) should contribute to investigations on rhythmic dysfunctions, such as epilepsy and schizophrenia. But in contrast to cortical microcircuitry, few detailed models exist for the thalamus (Hill and Tononi 2005; Iavarone et al. 2019; Izhikevich and Edelman 2008; Murray and Anticevic 2017).

In this study we present a NetPyNE implementation of the BBP S1 model, capturing most of the original single-cell physiology and morphology, synaptic mechanisms, connectivity and basic simulation results. With the addition of detailed thalamic circuits, we extend the results to show synchronous activity across cortical and thalamic populations, and open the door to new investigations on corticothalamic dynamics. The model is able to port readily across machines and can utilize a fast and efficient implementation on CPUs and GPUs using CoreNEURON. This extension allows the original BBP S1 model to be readily available to be used by the wider community to study a wide range of research questions.

## 2 Material and Methods

### 2.1 Individual neuron models

Cell reconstructions were based on the compartmental model Hodgkin-Huxley formalism, with membrane properties represented as components of an electric circuit, and ionic channels modeled as variable conductances. In the literature there are many large-scale brain models. (Shimoura et al. 2021) has discussed advantages on choosing each of those models and provided a guide on how to access connectivity maps and how to model brain inspired networks. To port the somatosensory microcircuit model in NetPyNE (Dura-Bernal et al. 2019), we recreated the single neuron models using cell files from the Neocortical Microcircuit Collaboration (NMCP; https://bbp.epfl.ch/nmc-portal) (Ramaswamy et al. 2015). The full dataset comprises 207 morpho-electrical (me) cell types, with 5 examples for each, totaling 1035 cell models, each stored with morphology file, descriptions of ion channels, and a NEURON HOC template to instantiate the cell, which can be imported directly to NetPyNE (Fig. 1). Neuron morphologies from the BBP S1 model (specifically, L1_DLAC, L4_DBC, L23_PC and L6_TPC_L4) imported into NetPyNE were visualized using the NetPyNE GUI (Fig. 1A). The full name of 207 cell types as well the corresponding acronym can be found in Table S1 in the Supplementary Material.

**Figure 1.**
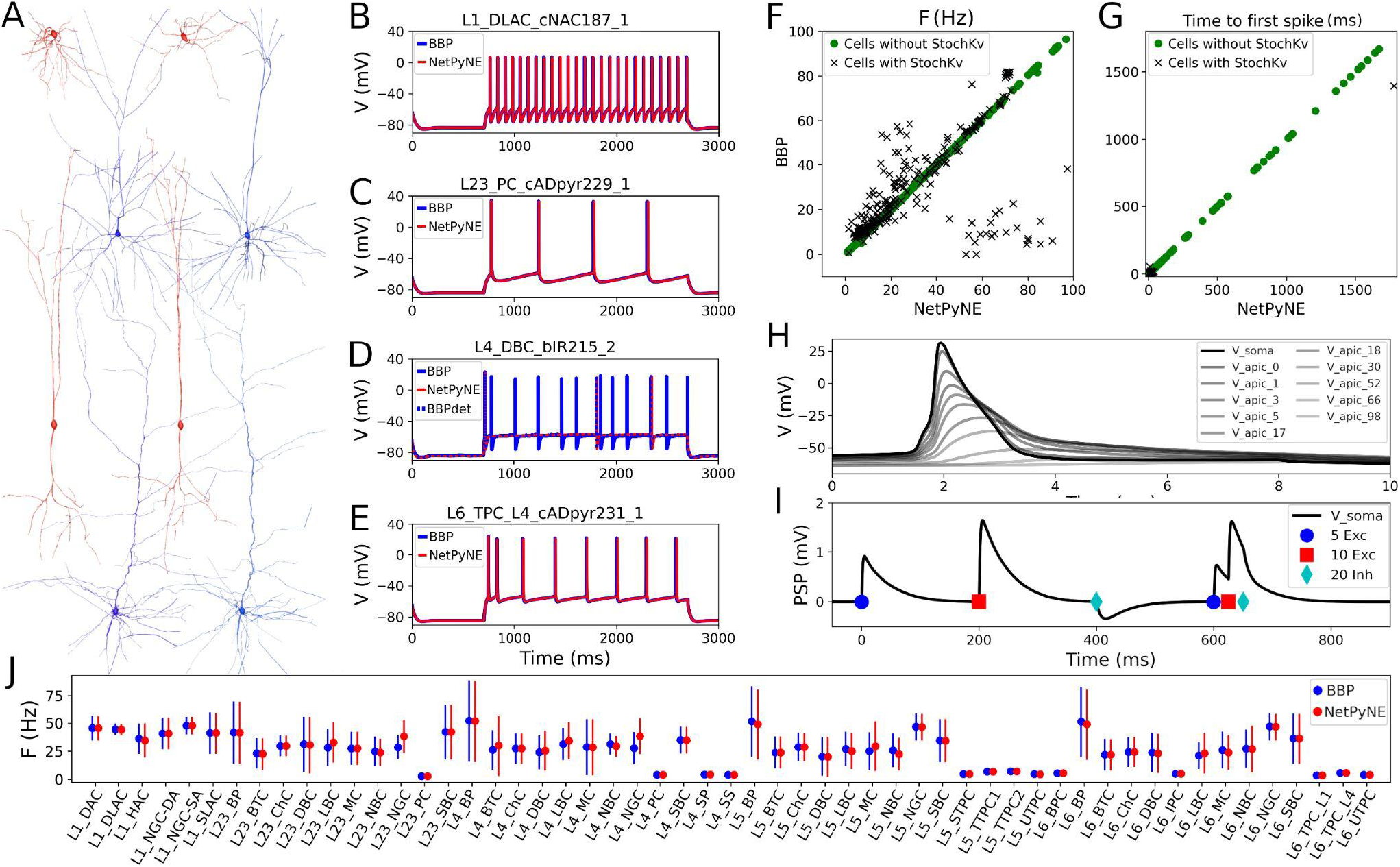
Reproduction and validation of BBP S1 cell types in NetPyNE. (A) 3D reconstructions of 4 pairs of m-type example neurons visualized using the NetPyNE graphical user interface: inhibitory cells L1_DLAC and L4_DBC (red), and excitatory cells L23_PC and L6_TPC_L4 (blue). (B-E) Somatic membrane potential of the neurons in (A) under current clamp with amplitude 120% of the neuron firing threshold. NetPyNE results (red) compared to the original BBP model results (blue). For L4_DBC_bIR cells (panel D) we used a deterministic version of the BBP stochastic potassium channel (StochKv) resulting in divergent results; using same deterministic channel in BBP (BBPdet, blue dotted line) restores the match to NetPyNE results. (F,G) Comparison of BBP and NetPyNE firing rate and time to first spike in response to current-clamp with amplitude 0.1 nA during 2 seconds for each cell type. Only some cell types with the StochKv show differences. (H) Backpropagating action potential in a L5_TTPC cell. (I) Post synaptic potentials (PSPs) of one dendritic connection with 5 excitatory (blue circles), 10 excitatory (red squares), and 20 inhibitory (cyan diamonds) synapses. (J) Comparison BBP and NetPyNE mean firing rate for all m-type populations. Due to the small number of cells with StochKv (3.6%), NetPyNE population firing rates closely match those of BBP.

Benchmark testing validated physiological responses (Fig. 1B-E) at 3 current clamp amplitudes (120%, 130%, and 140% of threshold; only 120% shown). Slight differences were observed in the cell types with a stochastic version of the K^+^ channel mechanism (StochKv; Fig. 1D) where we used a deterministic version of the channel from OpenSourceBrain (Gleeson et al. 2019; Gleeson et al. 2019b). The StochKv NMODL (.mod) mechanism required additional code outside of NetPyNE in order to update its state, and the inclusion of stochastic variables in each section of the cells significantly increased the simulation time. In order to understand StochKv effect on cell response, we applied a current clamp (0.1 nA, 2s) to the soma of each of the 1035 cells, and used the Electrophys Feature Extraction Library (eFEL) (eFEL 2015) to compare BBP and NetPyNE mean firing rate (Fig. 1F) and time to first spike (Fig. 1G) for those with and without the StochKv channel. As expected, variability with the StochKv channel in the original model was pronounced. Although present in 54/207 me-types, the StochKv channels are only in 3.63% of all cells. Within each m-type (morphology-type) those with StochKv also correspond to a minority of e-types (electrical) types; for example, only 32% of L4_DBC cells have e-type bIR (with StochKv channels). Given the small proportion of cells with StochKv channels (3.63%), the NetPyNE mean firing rates per m-type population closely match those of the original BBP model (Fig. 1J). Furthermore, the deterministic version of StochKv preserves irregular cell spiking patterns (CV_BBP_=0.25±0.16; CV_NetPyNE_=0.16±0.13; where CV is the inter spike interval coefficient of variation; see Supp. Fig. S1) as well as the neural firing rate (FR_BBP_=30.24±24.33, FR_NetPyNE_=28.67±21.20) in the current-clamp simulation with amplitude 0.1 nA during 2 seconds. For stimulation amplitude 0.8 nA, the CV (BBP=0.14±0.17; NetPyNE=0.15±0.30) and FR (BBP=115.58±70.20, NetPyNE=110.15±66.58) are similar in both model implementations (Supp. Fig. S1).

The NetPyNE implementation perfectly reproduced the original neuronal intrinsic dynamics since all model parameters were directly imported from the original HOC files, the same NMDOL files were used (except StochKv), and the underlying simulation engine was NEURON in both cases (see Figs. 1B-E). To validate, we simulated somatodendritic backpropagating action potentials (Fig. 1H) and dendrosomatic postsynaptic potentials (Fig. 1I) in an example L5_TTPC cell. Results were identical in the NetPyNE implementation and the original BBP cell models. To model dendrosomatic postsynaptic potentials (PSPs), we added excitatory connections with 5 and 10 synapses, and an inhibitory connection with 20 synapses, to the L5_TTPC neuron. Additionally, we provided the same three subthreshold inputs within a short time interval, to demonstrate temporal integration of PSPs (Fig. 1I).

### 2.2 Distribution and connectivity of cortical populations

Rather than instantiating the connectivity from a list of individual synapses based on anatomical overlap of neuronal arbors (M. W. Reimann et al. 2015), we created our S1 port using probability rules for both neuron distribution and connections. The network consisted of 31,346 cells in a cylindrical volume 2082 μm height and radius of 210 μm as in the original model (Fig. 2). Each population was randomly distributed within its specific layer (L1, L2/3, L4, L5, or L6). The number of cells in each one of 207 me-types was taken from the NMCP (Ramaswamy et al. 2015) the minicolumn data available was not used to distribute cells. A 2D representation of the cell distribution within the cylindrical volume is shown in Figure 2A, with layer thicknesses (in μm) for L1, L23, L4, L5, and L6 set to 165, 502, 190, 525, and 700, respectively. We used the S1 connectome (Gal et al. 2017) from NMCP, following the approach in (Michael W. Reimann, Horlemann, et al. 2017): 7 stochastic instances of a model microcircuit based on averaged measurements of neuron densities were used to calculate distance-dependent probabilities of connection. In each microcircuit instance, we calculated the connection probability for each pair of neurons based on the 2D somatic distance (horizontal XZ-plane) for each of the 1941 pathways. To estimate the distance-dependent probability, we calculated the probability in evenly spaced intervals starting at 25 ± 25 μm, in 50 μm intervals, up to 375 ± 25 μm. Next, we calculated the mean probability across the 7 microcircuits in evenly spaced intervals and used the mean values to fit the connection probability rules. We evaluated multiple functions for each pathway and selected the one that provided the best fit to the data. Figure 2B-D shows how this approach was used to calculate the connection probability of 3 example projections: data from the 7 microcircuit instances (mcs; cyan circles) is averaged across microcircuits (green diamonds) and fitted to either a single exponential (Fig. 2B, red line); an exponential with a linear saturation rule (Fig. 2C); or a single gaussian (Fig. 2D, dashed line). Because the original S1 model shows high variability in the number of synapses per connection, we calculated the mean values for each pathway and used it as a parameter in our model. The result is a representative reconstruction of the S1 column connectivity in NetPyNE, with approximately 27.6 million excitatory synapses and 9.6 million inhibitory synapses.

**Figure 2.**
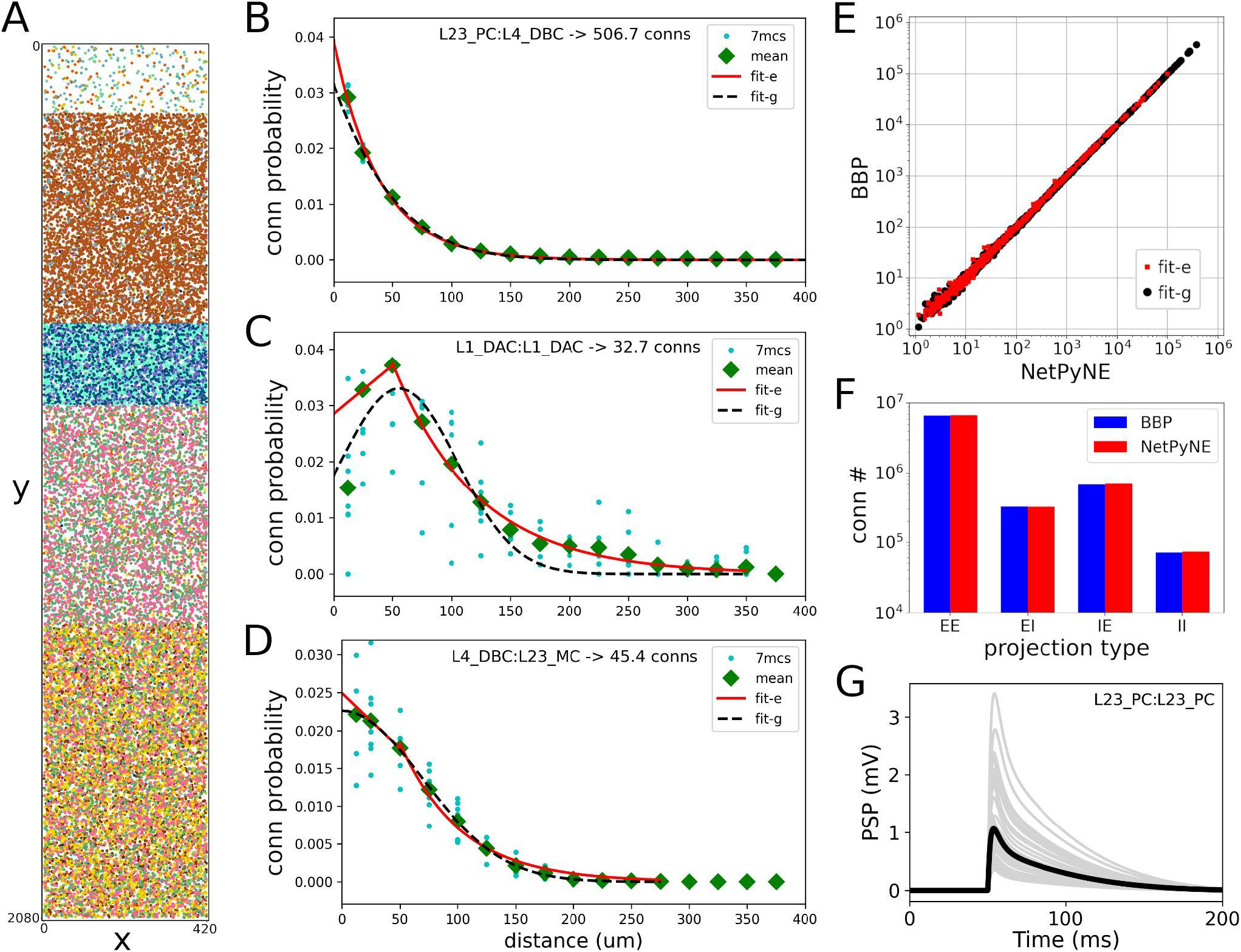
Reproduction and validation of BBP S1 neuron distribution and connectivity in NetPyNE. (A) 2D representation of the location of 31,346 cells in a cylinder with 2082 μm height and 210 μm radius, with each subtype (different colors) randomly distributed within its layer (L1, L2/3, L4, L5, or L6). (B-D) Probability of connection as a function of neuron pairwise 2D distance for three example pathways, each with a different best fit function: a single exponential (B, red line), exponential with a linear saturation rule (C, red line), and single gaussian fit (D, black dashed line). Cyan circles represent data from 7 microcircuit instances, and green diamonds represent the mean across the 7 instances. (E-F) Comparison of the number of connections between NetPyNE and BBP for each of the 1941 pathways (E) and 4 projection types (p-types) (F). (G) Postsynaptic potential (PSP) generated by connection between L23_PC neurons (Table 1, #18) simulated in NetPyNE; mean PSP trace (black line) across 20 PSP random instances (gray lines).

Using the fitted rules, we reconstructed an entire S1 column in NetPyNE and compared the two versions using the mean number of connections. To avoid overfitting, we generated 7 different instances using different connectivity seeds for both the NetPyNE and BBP models. The number of connections was similar in both models for each of the 1941 pathways (Fig. 2E) and for each of the four projection types (p-type) (EE, EI, IE, II) (Fig. 2F).

**Table 1.** Synaptic properties, s-type, p-type and rules for each class of connections implemented in NetPyNE. s-type: type of short-term dynamics; p-type: type of projection; g_syn_: peak conductance (ms); τ_decay_: decay time (ms); U: neurotransmitter release probability; D: time constant for recovery from depression (ms); F: time constant for recovery from facilitation (F). Values indicate mean ± standard deviation.

| # | BBP id | s-type | p-type | $g_{syn}$ (nS) | $\tau_{decay}$ (ms) | U | D (ms) | F (ms) | pre- and post-syn cell type rules |
| --- | --- | --- | --- | --- | --- | --- | --- | --- | --- |
| 0 | 0 | I1 | II | 0.83±0.55 | 10.40±6.10 | 0.16±0.100 | 45±21 | 376±253 | L6:L6_(DBC-LBC-NBC-SBC) |
| 1 | 3 | I1 | IE | 0.91±0.61 | 10.40±6.10 | 0.16±0.100 | 45±21 | 376±253 | SBC_cAC:Exc or L6_(NBC-LBC):L6_BPC |
| 2 | 13 | I1 | IE | 0.75±0.32 | 10.40±6.10 | 0.41±0.212 | 162±69 | 690±5 | L6_MC:L6_IPC |
| 3 | 1 | I2 | II | 0.83±0.55 | 8.30±2.20 | 0.25±0.130 | 706±405 | 21±9 | L1:Excitatory or Inhibitory:Inhibitory |
| 4 | 4 | I2 | IE | 0.91±0.61 | 8.30±2.20 | 0.25±0.130 | 706±405 | 21±9 | SBC_dNAC:Excitatory |
| 5 | 8 | I2 | IE | 0.75±0.32 | 8.30±2.20 | 0.25±0.130 | 706±405 | 21±9 | BTC-DBC-BP:Excitatory |
| 6 | 9 | I2 | IE | 0.75±0.32 | 8.30±2.20 | 0.30±0.080 | 1250±520 | 2±4 | MC:Excitatory |
| 7 | 10 | I2 | IE | 0.91±0.61 | 8.30±2.20 | 0.14±0.050 | 875±285 | 22±5 | LBC-NBC_(bAC cAC bNAC dNAC):Excitatory |
| 8 | 12 | I2 | IE | 2.97±0.95 | 8.30±2.20 | 0.25±0.130 | 706±405 | 21±9 | Chc:Excitatory |
| 9 | 5 | I3 | IE | 0.91±0.61 | 6.44±1.70 | 0.32±0.140 | 144±80 | 62±31 | SBC_bNAC or LBC-NBC_(cNAC dSTUT cSTUT bSTUT):Excitatory |
| 10 | 11 | I3 | IE | 0.83±0.55 | 36.55±0.71 | 0.25±0.130 | 706±405 | 21±9 | NGC:Excitatory |
| 11 | 114 | E1 | EI | 0.43±0.28 | 1.74±0.18 | 0.02±0.001 | 194±10 | 507±20 | Exc:(BP_cAC DBC_cAC BTC_cAC) |
| 12 | 115 | E1 | EI | 0.72±0.50 | 1.74±0.18 | 0.02±0.001 | 194±10 | 507±20 | Exc:(NBC-LBC)_(cAC cIR bAC bIR cNAC) |
| 13 | 132 | E1 | EI | 0.72±0.50 | 1.74±0.18 | 0.01±0.001 | 242±15 | 563±32 | L6_TPC_L:L6_(DBC-LBC-NBC-SBC) |
| 14 | 133 | E1 | EI | 0.11±0.08 | 1.74±0.18 | 0.09±0.120 | 138±211 | 670±830 | Excitatory:MC |
| 15 | 116 | E2 | EE | 0.72±0.50 | 1.74±0.18 | 0.50±0.020 | 671±17 | 17±5 | Excitatory:Excitatory |
| 16 | 117 | E2 | EI | 0.43±0.28 | 1.74±0.18 | 0.50±0.020 | 671±17 | 17±5 | Excitatory:[L1-BP_(cNAC bNAC)-DBC_bAC-BTC_(bAC cNAC bIR)] |
| 17 | 118 | E2 | EI | 0.72±0.50 | 1.74±0.18 | 0.50±0.020 | 671±17 | 17±5 | Excitatory:SBC-ChC |
| 18 | 119 | E2 | EE | 0.68±0.46 | 1.74±0.18 | 0.46±0.260 | 671±17 | 17±5 | L23_PC:L23_PC |
| 19 | 120 | E2 | EE | 0.68±0.46 | 1.74±0.18 | 0.86±0.049 | 671±17 | 17±5 | L4_Excitatory:L4_Excitatory |
| 20 | 121 | E2 | EE | 0.19±0.12 | 1.74±0.18 | 0.79±0.040 | 671±17 | 17±5 | L4_SS:L23_PC |
| 21 | 122 | E2 | EE | 0.80±0.53 | 1.74±0.18 | 0.39±0.030 | 671±17 | 17±5 | L5_STPC:L5_STPC |
| 22 | 123 | E2 | EE | 1.50±1.05 | 1.74±0.18 | 0.50±0.020 | 671±17 | 17±5 | L5_TTPC:L5_TTPC |
| 23 | 127 | E2 | EE | 0.80±0.53 | 1.74±0.18 | 0.39±0.134 | 780±54 | 51±36 | L6_IPC:L6_IPC |
| 24 | 131 | E2 | EI | 0.72±0.50 | 1.74±0.18 | 0.58±0.070 | 240±43 | 71±47 | L6_IPC:L6_(DBC-LBC-NBC-SBC) |
| 25 | 134 | E2 | EI | 0.72±0.50 | 1.74±0.18 | 0.72±0.065 | 227±38 | 14±12 | Exc:(NBC-LBC)_(bSTUT dNAC bNAC cSTUT) |
| 26 | 126 | E3 | EE | 0.80±0.53 | 1.74±0.18 | 0.21±0.032 | 460±53 | 230±69 | L6_TPC_L:L6_TPC_L |
| 27 | 128 | E3 | EE | 0.80±0.53 | 1.74±0.18 | 0.27±0.033 | 559±238 | 200±92 | L6_IPC:L6_BPC |
| 28 | 129 | E3 | EE | 0.80±0.53 | 1.74±0.18 | 0.22±0.053 | 535±134 | 116±81 | L6_IPC:L6_TPC_L |

### 2.3 Synaptic physiology

The original BBP S1 model included detailed synaptic properties (conductances, post-synaptic potentials, latencies, rise and decay times, failures, release probabilities, etc) recapitulating published experimental data. Short-term dynamics were used to classify synapses into the following types (s-types): inhibitory facilitating (I1), inhibitory depressing (I2), inhibitory pseudo-linear (I3), excitatory facilitating (E1), excitatory depressing (E2), and excitatory pseudo-linear (E3). A set of rules were then derived from experimental data to assign an s-type to each broad class of connections. Based on the NMCP data, there were 29 classes of connections as determined by the combination of pre- and post-synaptic me-types. The synaptic properties, s-type and p-type for each class of connections are summarized in Table 1.

The dual-exponential conductance model with rise time (τ_rise_) 0.2 ms was used for all synapses. Moreover, synaptic properties included the kinetic parameters: peak conductance (g_syn_) and decay time (τ_decay_); and dynamic parameters: neurotransmitter release probability (U), time constant for recovery from depression (D; in ms) and time constant for recovery from facilitation (F; in ms). The NetPyNE implementation reproduces the original PSP amplitudes from Markram et al (2015). An example of PSPs for a connection between L23_PC neurons (Table 1, #18) simulated in NetPyNE is shown in Fig. 2G. The mean PSP peak amplitude across 20 PSPs (with different randomization seeds) was 1.0 mV, which matches the value obtained in Markram et al. (2015). We also included a compact description of the rules to determine what connections belong to each class, based on the pre- and postsynaptic cell types (Table 1). For clarity, we rearranged the classes of connections by s-types in the sequence I1, I2, I3, E1, E2, and E3 (from 0 to 28), and included the original BBP class label for reference. The parameters D and F correspond to the synapses with short term plasticity (STP), which could be optionally added to recurrent S1 connections, and connections from thalamus to S1.

The s-types for each class of connections and for each of the 1941 pathways are color-coded and illustrated in Figure 3A. Since pathways depend on m-types but connection classes depend on me-types (each m-type includes multiple me-types), it is possible to have multiple s-types for the same pathway, in those cases we simply labeled it as either I2 or E2. To implement the dynamics of each s-type in NetPyNE we used a deterministic version of the dual-exponential synaptic model (Hennig 2013; Fuhrmann et al. 2002). Example simulations of the post-synaptic potentials for the different s-types are shown in Figure 3B. For each example, we ran 20 simulations with 5 different post-synaptic cells of the same me-type and 4 random synaptic distributions. Pre- and post-synaptic neurons of specific me-types were selected to illustrate each of the six s-types (see Fig. 3 caption).

**Figure 3.**
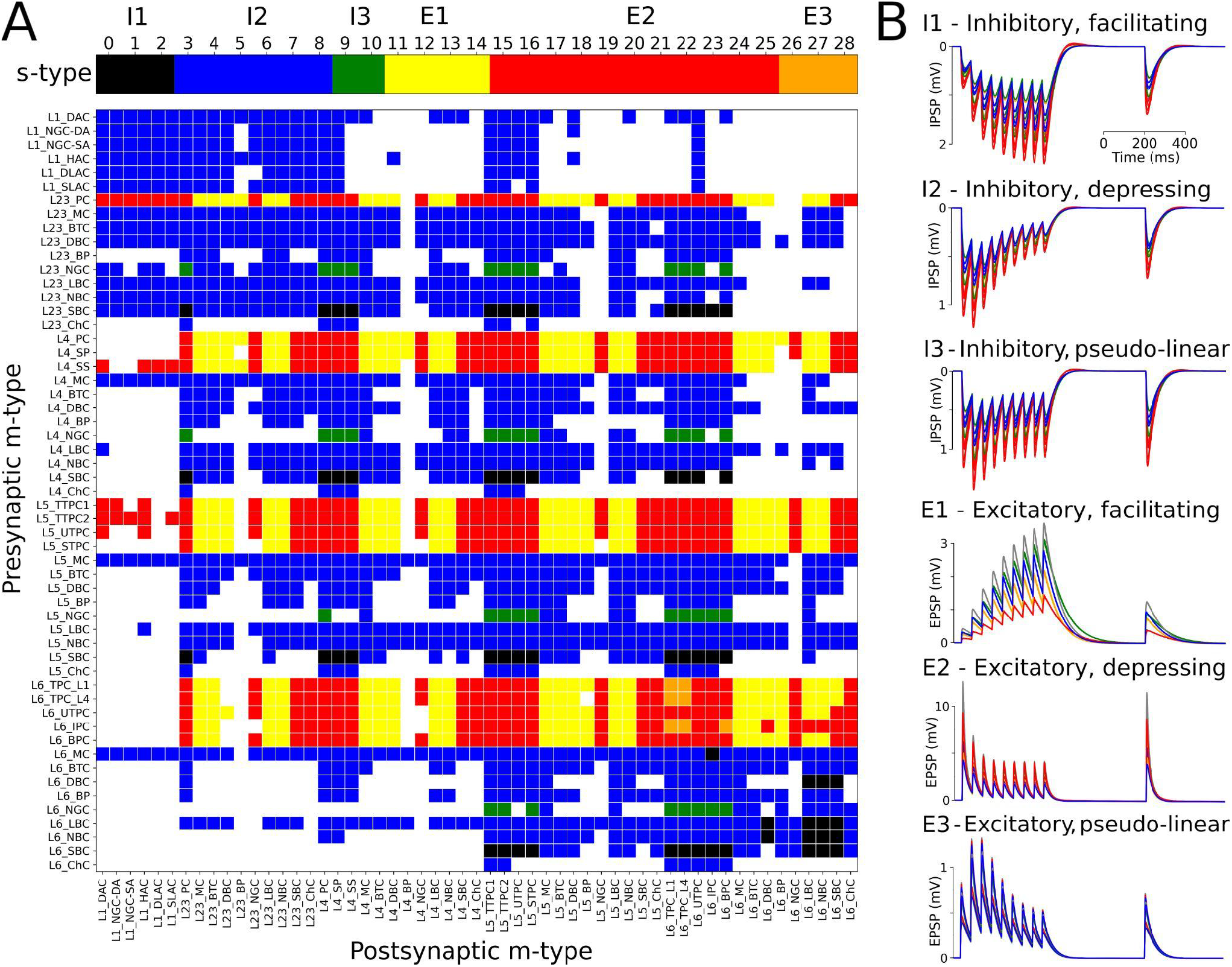
Matrix of s-types for each of the 1941 pathways and simulated PSPs exemples for each s-type in NetPyNE. (A) Color-coded s-types for each class of connections (top) and for each of the 1941 pathways (bottom). Note that for pathways with multiple s-types only either I2 or E2 is shown. (B) Example simulations of post-synaptic potentials to illustrate each of the six s-types. Each example shows the results of 20 simulations with 5 different post-synaptic cells (different colors) of the same me-type, and 4 random synaptic distributions. Inhibitory s-types were simulated using pathway L23_SBC:L23_PC, which included different s-types depending on pre-synaptic e-type: I1 for e-type cAC, I2 for e-type dNAC, and I3 for e-type bNAC (as shown in Table 1). Excitatory s-types E1 and E2 were simulated using pathway L23_PC:L23_LBC, e-types cAC and dNAC, respectively; and s-type E3 was simulated using pathway L6_TPC_L4:L6_TPC_L4.

### 2.4 Extending the model to include thalamic populations and connectivity

We extended the model to include somatosensory thalamic populations with cell type-specific dynamics, intra-thalamic connectivity and bidirectional projections with cortex. In the original model, thalamic inputs were modeled as spike generators that only provided feedforward inputs to S1. Our somatosensory thalamus model is composed of the excitatory ventral posterolateral (VPL), ventral posteromedial (VPM) and the posteromedial (POm) nuclei, and the inhibitory reticular nucleus (RTN). We used single compartment cell models with dynamics tuned to reproduce previous studies on the interaction between the thalamic relay and reticular cells (Destexhe, Bal, et al. 1996), but adjusted to work in large-scale networks (Moreira et al. 2021). The thalamic circuit architecture consisted of six stacked populations as a rough approximation of the thalamic anatomical layout (Fig. 4A). The top three were inhibitory populations comprising the outer, middle and inner sectors of the RTN, and spanning a height of 78 μm, 78 μm and 156 μm, respectively. Below these were the three excitatory populations, VPL, VPM and POm, with heights of 156 μm, 156 μm and 312 μm, respectively. The horizontal dimensions (XZ-plane) for all populations were 420 μm x 420 μm. Cells were randomly distributed across each nuclei with the number of cells in each population based on cellular density obtained from the Cell Atlas for the Mouse Brain (https://bbp.epfl.ch/nexus/cell-atlas/) (Erö et al. 2018). Although POm was larger than VPL and VPM, we reduced its cell density by 50%, resulting in approximately the same population size. This lower density accounts for the proportion of coexisting, but functionally-isolated, M1-projecting thalamocortical cells present in POm with no projections to S1 (Guo et al. 2020).

**Figure 4.**
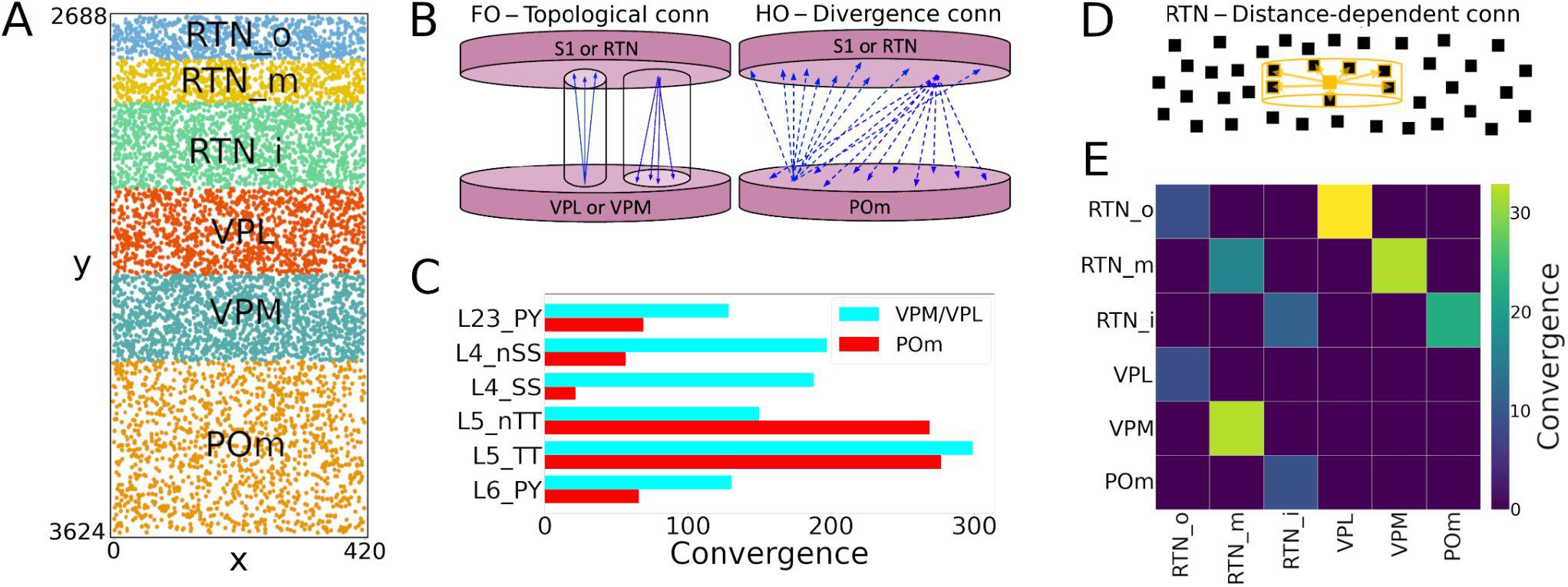
NetPyNE model of somatosensory thalamic populations and connectivity extending the original S1 model. (A) Distribution of neurons across the six different thalamic populations, roughly mimicking the thalamus anatomy (x and y axes in μm). (B) Schematic of bidirectional connectivity between thalamic regions and cortex. Bidirectional connections between S1/RTN and first order (FO) regions VPL and VPM are topological, whereas those with high order (HO) region POm are non-topological and implemented using divergence rules. (C) Convergence connectivity between thalamic regions and S1 (D) RTN cells have sector-specific distance-dependent connectivity (E) Convergence connectivity matrix across all thalamic populations.

The intrathalamic connectivity was based on data of axonal and dendritic footprints for each nucleus. The VPL and VPM are considered first-order nuclei (FO), which means they receive afferent information from peripheral sensory organs (not modeled here) and are interconnected with cortex (Ma 1991; Sugitani et al. 1990; Łuczyńska et al. 2003) and RTN (Lam and Sherman 2011) in a topological fashion. On the other hand, POm is considered a higher-order (HO) nucleus, so input arrives mainly from the cortex, in this case, from S1 L5 and L6 (O’Reilly et al. 2021; Ohno et al. 2012). The connectivity pattern of HO nuclei has not been properly characterized, but literature reports a decreased level of organization of HO nuclei inputs to RTN (Lam and Sherman 2011), as it sends projections to S1.

We therefore adopted three connectivity strategies. In the first, neurons from FO nuclei projected with a column-like topological organization. We implemented this by combining a probability of connection that decreased exponentially with the horizontal distance between the pre- and post-synaptic cells with a decay constant proportional to the footprint radius, and which was truncated to 0 outside of the footprint radius (this denotes the maximum distance of connection in the XZ-plane). The following footprint diameters were derived from experimental data (or estimated in the case of no literature reports) for the different axonal footprints of each thalamic projection: RTN→VPL and RTN→VPM: 64.33 μm (Lam and Sherman 2007); VPL→RTN: 97.67 μm; and VPM→RTN: 103.57 μm (Lam and Sherman 2011). The second strategy applies to RTN→RTN connectivity and implements a sector-specific distance-dependent connectivity. More specifically, within each RTN sector, the probability of connection decayed exponentially and was truncated to 0 based on a footprint radius of 264.63 μm (Lam, Nelson, and Sherman 2006). In strategies one and three, the maximum distance in the Y-plane was set to 10% of the footprint radius, following the disc-like morphology from the axonal projections of the relay cells and the dendritic trees of reticular cells (Lam, Nelson, and Sherman 2006; Murray Sherman and Guillery 2001). The third strategy was a divergence rule, with the number of projections from and to HO nuclei having a fixed value and being distributed without spatial constraints. This divergence value was adjusted so that the HO dynamics resembled that of the FO nuclei. This allowed us to replicate a column-like topological organization in FO nuclei using single-compartment cells (Lam and Sherman 2007), and distribute the HO connections to behave in a functionally similar fashion (Fig. 4B).

All excitatory thalamic nuclei were indirectly interconnected through their RTN projections, which was divided into three sectors, in line with reports of preferred innervation zones by each of the thalamic nuclei (Lam and Sherman 2011). Synapses within RTN were mediated by GABA_A_, from RTN to the excitatory nuclei by a combination of GABAa and GABAb with equal weight, and by AMPA from the excitatory nuclei to RTN and cortex (Destexhe, Bal, et al. 1996). The probability and weight of connections were the targets of parameter optimization. The matrix with the convergence of intra-thalamic connections is shown in Figure 4E.

Feedback corticothalamic connectivity originated from S1 m-types L5_TTPC2 and L6_TPC_L4 (O’Reilly et al. 2021). Similar to the topological rules described above, we implemented connectivity with convergence of 30 (i.e. number of pre-synaptic cells projecting to each post-synaptic cell), but only if the horizontal distance between the pre- and post-synaptic neurons was lower than 50.0 μm (Fig. 4).

Thalamocortical connectivity from VPL, POm and VPM to S1 was implemented using convergence values estimated from previous studies (Meyer et al. 2010) (Fig. 4C). Convergence values for each of the 55 m-types were calculated based on the weighted average of the populations in each layer. The convergence values for inhibitory populations were multiplied by a scaling factor derived from the original mode (∼0.595). This thalamic convergence factor for inhibitory cells was estimated by dividing the IE ratio of VPM thalamic innervation (83/775 = 0.107) by the average IE population ratio (4,779/26,567 = 0.18). The resulting S1 column received approximately 4.95M synapses from VPM, 4.95M from VPL, and 3.1M from POm. This is consistent with values that can be derived from experimental studies (Meyer et al. 2010), with 4.27M synapses from VPM and, and 2.66M from POm. We approximated thalamocortical (TC) synaptic physiology using the parameters of model #25 in Table 1, and using 9 synapses per connection, following Markram et al. (2015) characterization of TC synapses as excitatory depressing (E2).

### 2.5 Background inputs

Each cell in the S1 circuit received 10 synaptic inputs from Poisson-distributed spike generators (NetStims) to represent the global effect of spontaneous synapses, background, and other noise sources from non-modeled brain regions projecting to S1. These stimuli were randomly distributed randomly across all sections. The quantal synaptic conductance was calculated based on the average quantal conductance for excitatory and inhibitory synapses. We tuned the excitatory and inhibitory stimuli rates using grid search parameter exploration to obtain average excitatory firing rates of ∼1Hz and physiological firing rates for most S1 populations.

### 2.6 Model building

We used the NetPyNE modeling tool (Dura-Bernal et al. 2019) to build, manage simulations, and analyze results of the S1 and thalamic circuit model. NetPyNE employs NEURON (Lytton et al. 2016; Carnevale and Hines 2006; Migliore et al. 2006) as backend simulation engine, with either the standard or CoreNEURON libraries (Kumbhar et al. 2019). The high-level Python-based declarative language provided by NetPyNE facilitated the development of this highly complex and extensive circuit model. This language enabled us to easily import existing morphological and biophysical parameters of different cell types, and define complex connectivity and stimulation rules. We used NetPyNE to explore and optimize the model parameters through automated submission and managing of simulations on supercomputers. We also employed NetPyNE’s built-in analysis functions to plot 2D representations of cell locations, connectivity matrices, voltage traces, raster plots, local field potentials (LFPs), 3D synapses representations, and firing rate statistics. NetPyNE can also be used to export the model into the NeuroML (Gleeson et al. 2010) and SONATA (Dai et al. 2020) standard formats.

Model parameters are based on experimental data and the original model (Markram et al. 2015). Nonetheless, parameter optimization was necessary to ensure the model reproduces experimental measures such as population firing rates and post-synaptic potentials (PSP). The parameters optimized for the S1 thalamocortical circuit were the background rate for excitatory and inhibitory connections. For the intrathalamic projections, we optimized connection weight (range 0 to 2 mV), connection probability (range 0 to 1), y-axis connection radius (1, 2, 5 or 10%) and connectivity divergence of the HO populations (5, 10, 20 or 40 cells). For the thalamocortical and corticothalamic projections we optimized connection weight (range 0 to 2 mV) and connection probability (range 0 to 1). More details about model parameter optimization/exploration are described in Section 2 of the Supplementary Material.

### 2.7 Simulation of local field potentials (LFPs)

We simulated LFP extracellular recordings using the “line source approximation” (Buzsáki, Anastassiou, and Koch 2012; Łęski et al. 2013; Parasuram et al. 2016), which is based on the sum of the transmembrane currents generated by each segment of each neuron, divided by the distance between the segment and the electrode. This method assumes that the electric conductivity (sigma = 0.3 mS/mm) and permittivity of the extracellular medium are constant everywhere and do not depend on frequency. LFP calculation, analysis and visualization was performed using NetPyNE.

Given the computational cost and memory requirements of simulating the full S1 model with morphologically-detailed neurons while recording LFPs, we calculated transmembrane currents only for the most central cells within an 84 μm (20% of 420μm) diameter cylinder. This means that only 4.4% of the neurons were simulated in full detail, i.e. using full morphological reconstructions and with all synapses from the full model. The dynamics of the remaining cells (∼96% S1 and thalamus) were simulated using spike generators (VecStims in NEURON) using the spiking activity previously recorded in full simulations. That is, the inputs and activity of the 4% of fully detailed neurons from which the LFPs were calculated were identical to those of the full network simulations (when 100% of the neurons are simulated in detail).

## 3 Results

### 3.1 Reproduction of cell morphologies, physiological responses, spatial distribution and connectivity

Cells imported into NePyNE using the files from The Neocortical Microcircuit Collaboration NMCP (Ramaswamy et al. 2015), reproduced the morphological and electrophysiological characteristics of the original model (Fig. 1): mean firing rate and time to the first spike after a current clamp stimulation are fitted for all the 1035 cell types. Firing dynamic differences were observed in cells with the stochastic K channel (StochKv), but their firing irregularity was partly preserved (Supp. Fig. S1) and, given their low proportion (3.63%), the average firing rates of all m-type populations closely matched those in the original model (Fig. 1H).

We were able to recreate the general characteristics across the 7 BBP S1 microcircuit instances: the 31,346 cells distributed randomly by layer, and probabilistic connections were generated for each of the 1941 pathways (Fig. 2). Here, we replaced the original connectivity method, based on the overlap between axonal and dendritic fields, with one based on connection probability based on cell type, layer, inter-cell distance, and dendritic pattern of post-synaptic locations. This network parameterization allowed us to rescale the microcolumn and generate different instances by changing the random number generator seed. Our probabilistic rules best reproduced the original number of connections using a Gaussian fit in most projection pathways (1303 of 1941) and an exponential fit plus a linear saturation in the remaining 638 cases (Fig. 2).

### 3.2 Extension to include detailed thalamic circuits

We extended the model to include the somatosensory thalamic populations with projections to S1: RTN, POm, VPL, and VPM. The number of thalamic cells was adapted to fit a cylindrical column with the same radius as the S1 column. This facilitated the inclusion of topological connectivity rules between the two regions. We reproduced the firing dynamics of the different thalamic cell types using a single compartment neuron model (Moreira et al. 2021). The connections from TC cells to S1 were based on convergence rules derived from experimental data (Meyer et al. 2010), and synaptic physiological mechanisms were generalized from the BBP VPM projections to S1 layers 4 and 5 (Markram et al. 2015). Feedback connections originated from S1 cell types L5_TTPC2 and L6_TPC_L4 and targeted VPL and VPM following a topological organization, and POm in a following a non-topological broader distribution (Fig. 4). The parameters of the thalamic circuit were adjusted to reproduce a stable self-sustained activity with rhythmic bursting and spindle oscillations (Destexhe and Contreras 2011), as well as a shift in dynamics following localized excitatory input in the relay cells (Bonjean et al. 2012; Moreira et al. 2021).

### 3.3 Cortical and thalamic circuits independent response to background inputs (no thalamocortical connections)

We first evaluated the response to background inputs of the S1 cortical circuit and the thalamus circuit independently, i.e., without any connections between cortex and thalamus. When driven with background inputs the S1 model generated spontaneous activity with most populations (48 out of 55) firing within physiological rates (Fig. 5). To achieve this, excitatory and inhibitory background inputs were tuned via grid search parameter optimization (see Methods). Figure 5 illustrates the S1 spontaneous activity results, including a spiking raster plot of all 31,346 cortical cells, examples of voltage traces for each of the 207 me-type population, and the average firing rates for each of the 55 m-type populations. The thalamic populations, disconnected from S1 and driven by background inputs, exhibited stable self-sustained activity with rhythmic bursting at theta ∼6 Hz (Kim and McCormick 1998) (Fig. 5C,D). These oscillations, which were most prominent in the RTNi and POm populations, emerged despite the lack of rhythmicity in the background inputs. The thalamic circuit oscillatory dynamics are consistent with the recurrent interactions between thalamic relay and reticular neurons described in previous studies (Destexhe, Bal, et al. 1996)

**Figure 5.**
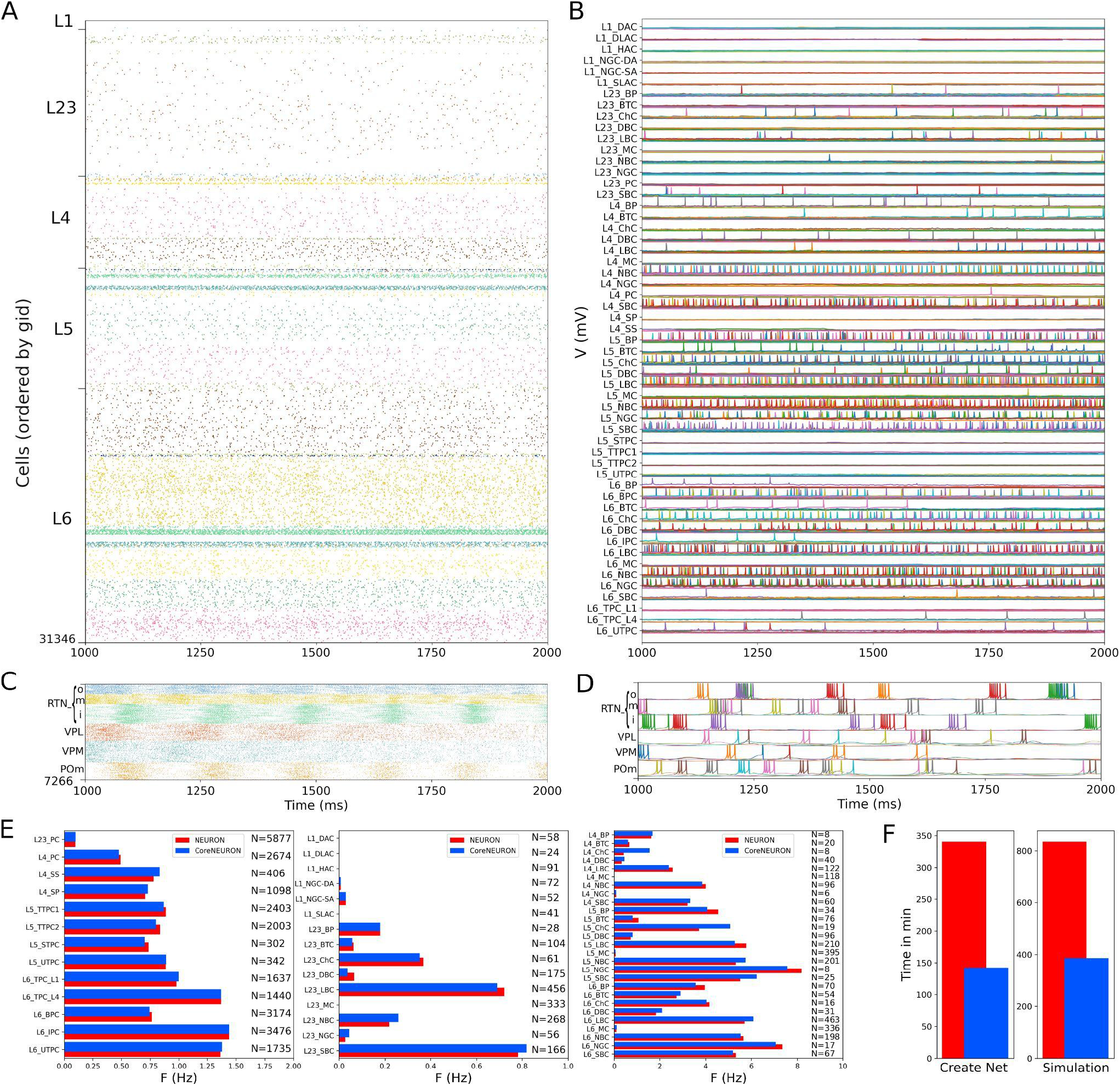
NetPyNE S1 and thalamus circuit response to background inputs (spontaneous activity). (A) Spiking raster plot of the 31,346 cells in the S1 column during 1 second (the first second was omitted to allow the network to reach a steady state). (B) Example voltage traces for each of the 207 me-types grouped by rows into their respective 55 m-types (same time period as raster plot). (C) Spiking raster plot of the 7,266 thalamic neurons during 1 second showing intrinsic oscillations. (D) Example voltage traces for each of the 6 thalamic populations. (E) Mean firing rates of each of the 55 m-types for NEURON (red) vs CoreNEURON (blue). (F) Comparison of the time required to create the network and run the simulation on a 40-core Google Cloud virtual machine using NEURON (red) or CoreNEURON (blue).

Simulations were run using NetPyNE and NEURON on a Google Cloud virtual machine with 40 cores. We compared the S1 results using the standard NEURON simulation engine vs CoreNEURON, a state-of-the-art solver optimized for large scale parallel simulations on both CPUs and GPUs (Kumbhar et al. 2019). Both simulation engines produced very similar firing rates for each population (Fig. 5E), with excitatory and L1-L3 inhibitory cells showing overall lower firing rates than L4-L6 inhibitory cells. The overall average firing rate across the 2 simulated seconds was 0.95 Hz in both cases (NEURON: 59,779 spikes; CoreNEURON: 59,749 spikes). This demonstrates the consistency of results obtained from both simulation engines, making CoreNEURON a viable alternative to study the S1 network. CoreNEURON was 2.4x faster to create the network and 2.2x faster to run the simulation (Fig. 5F).

### 3.3 S1 circuit response to background inputs with short term plasticity (no thalamocortical connections)

We simulated the response of the S1 cortical circuit to background inputs but including short term plasticity (STP) in its local synaptic connections (Fig. 6A). Adding STP resulted in the emergence of synchronous bursting within the S1 cortical column at approximately 1 Hz frequency (compare S1 raster in Figs. 5A and 6A). The spontaneous synchronous bursts first appeared in L5, and then spread to all S1 cells within 100 ms. Figure 6B shows an amplified raster plot of L4-L6 with 70 ms of activity at the time when spontaneous synchronous bursts started. Figure 6C shows example voltage traces of cortical and thalamic neurons, illustrating the spike synchrony of S1 and the thalamic bursts. These results are comparable to the simulations presented in Figs. 11B,C of the original publication (Markram et al. 2015).

**Figure 6.**
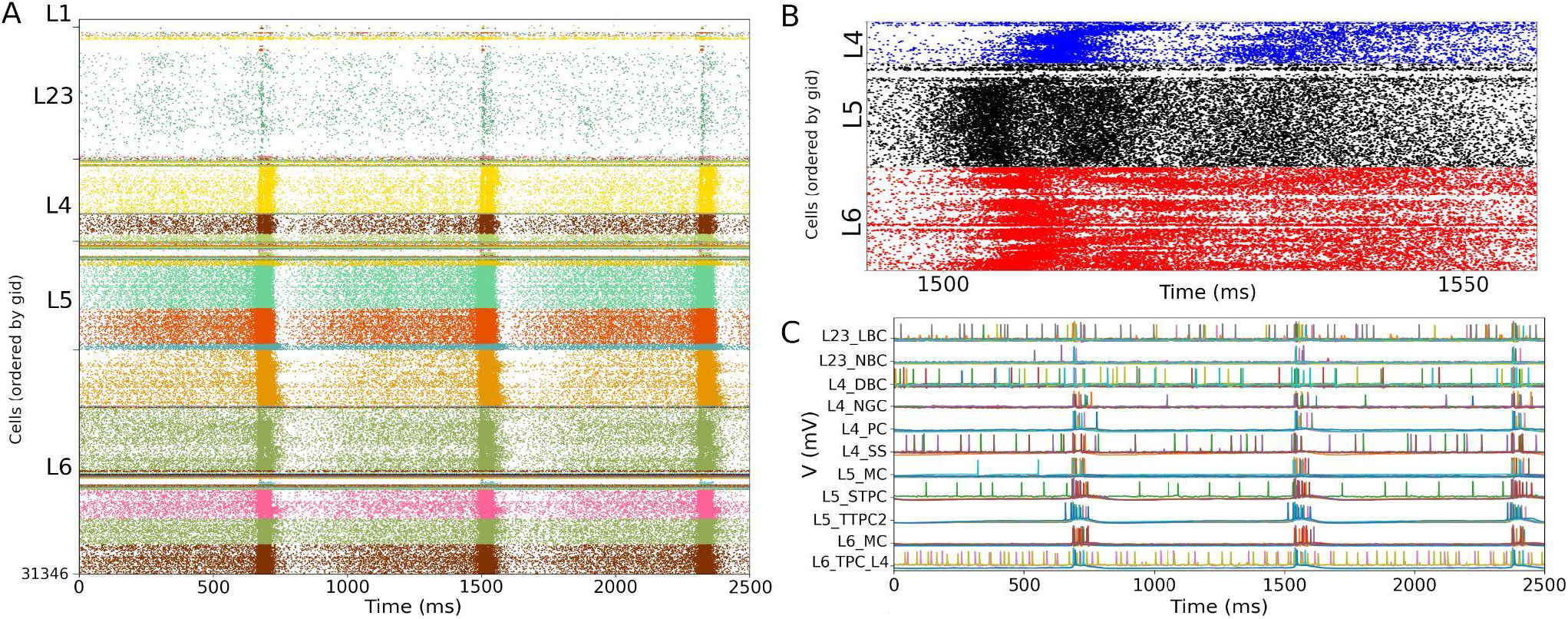
NetPyNE S1 circuit response to background inputs with short-term plasticity (STP). (A) Spiking raster plot of S1 with STP. S1 and thalamus were not interconnected; only intracortical connections were included. (B) Amplified spiking raster plot (A) showing the 70 ms around the time when synchronous bursts first occur in L5 (black) and then propagate to L6 (red) and L4 (blue). (C) Example traces from (A) showing spike synchrony across cortical populations and thalamic bursts. Rasters in A show 2.5 seconds after steady state was reached.

### 3.4 S1 and thalamic circuit response with bidirectional thalamic connectivity and cortical short term plasticity

We then simulated the full circuit with bidirectional connections between S1 and thalamus and STP in the thalamus to S1 connections (Fig. 7A). The full cortico-thalamo-cortical circuit exhibited overall increased activity with S1 oscillations around 6 Hz frequency, and strong thalamic oscillatory activity at the same frequency. Oscillations were now synchronized across all S1 and thalamic populations. Figure 7B shows the voltage traces of several cortical and thalamic neurons, illustrating the spike synchrony of S1 and thalamic populations. Finally, in Figure 7C we compare the mean firing rate for all S1 and thalamic populations with (red bars) and without (blue bars) bidirectional thalamocortical connectivity. All 55 model populations now exhibited physiological firing rates. Adding bidirectional thalamocortical connectivity resulted in a modest increase of the overall mean firing rate, from 4.96 Hz to 5.29 Hz, with more pronounced increases in the average firing rates of L1 and L2/3 inhibitory populations. These results do not have a direct correspondence to any in the original BBP publication, since the original model did not include thalamic populations bidirectionally connected to cortex.

**Figure 7.**
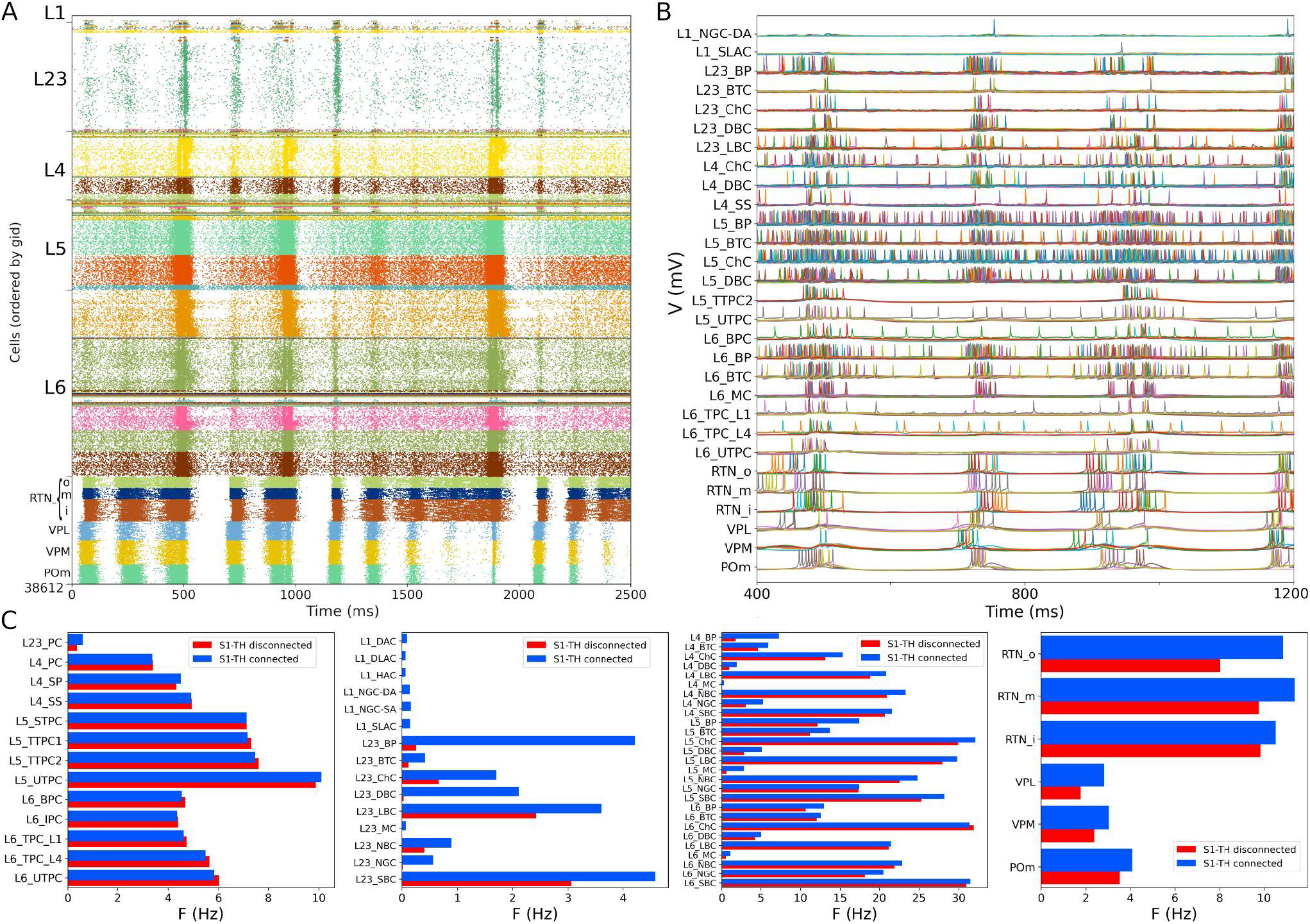
NetPyNE S1 and thalamic circuit response with bidirectional thalamic connectivity and cortical STP. (A) Spiking raster plot of the fully connected circuit model, including bidirectional connections between S1 and thalamus (shows 2.5 seconds of simulation after steady state was reached). Oscillations at ∼6 Hz were now synchronized across all S1 and thalamic populations. (B) Example traces from (A) during 800 ms showing spike synchrony across cortical and thalamic populations. (C) Comparison of mean firing rates of each of the 55 S1 and 6 thalamic m-types with (*S1-TH connecte*d) and without (*S1-TH disconnected*) bidirectional thalamocortical connectivity (compare rasters in Figs. 7A and 6A, respectively).

### 3.5 S1 and thalamic circuit response after reducing the extracellular calcium concentration to reproduce asynchronous in vivo-like state

Experimental evidence shows that extracellular calcium concentration ([Ca^2+^]_o_) in vivo is lower than in vitro, and, as a consequence, PSP amplitudes are also lower (Borst 2010). Markram et al. (2015) divided the dependency of PSPs on [Ca^2+^]_o_ into three classes for specific connection types: steep, intermediate, and shallow. The PSP amplitudes have steep dependence for connections between PC-PC and PC-distal targeting cell types (DBC, BTC, MC, BP) and a shallow dependence for connections between PC-proximal targeting (LBCs, NBCs, SBCs, ChC). An intermediate level of dependence was assumed for other connections. To simulate reduced [Ca^2+^]_o_ in the NetPyNE implementation we decreased the *cao* parameter from 2.0 to 1.2 in all cells, and modified the use parameter of synaptic transmission (U) adding a factor to multiply its value from 0.25 to 0.75. This resulted in a transition from synchrony (in vitro-like, Figs. 6,7) to asynchrony (in vivo-like, Fig. 8) network states, as in Markram et al. (2015). The increased asynchrony happened both for the S1-TH disconnected (Fig. 8A) and the S1-TH connected (Fig. 8B) cases. Decreasing extracellular calcium concentration resulted in decreased firing rates for most populations (compare Figs. 7C and 8C). In both the in vitro and in vivo conditions, cortical firing rates were generally slightly higher for the S1-TH connected case. However, bidirectional thalamic connectivity (S1-TH connected) resulted in increased thalamic population firing rates in vitro, whereas under in vivo conditions (low [Ca^2+^]_o_), it lowered thalamic firing rates and decreased synchrony.

**Figure 8.**
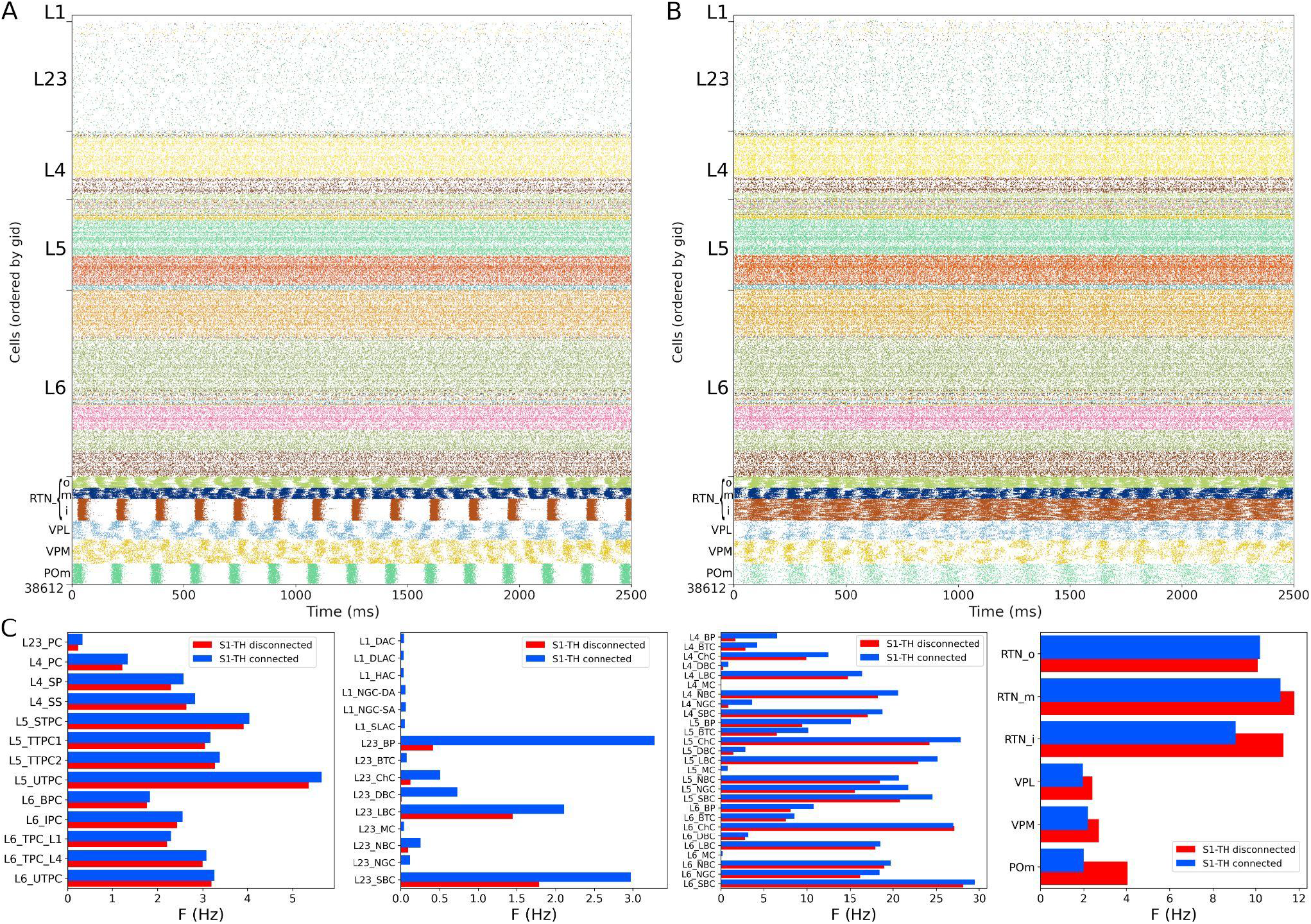
NetPyNE S1 and thalamic circuit response with low extracellular calcium (in vivo-like asynchronous states). (A) Spiking raster plot of spontaneous activity with only intracortical and intrathalamic connections (*S1-TH disconnected*). (B) Spiking raster plot of the fully connected circuit model, including bidirectional connections between S1 and thalamus (*S1-TH connected*). (C) Comparison of mean firing rates of each of the 55 S1 and 6 thalamic m-types without (*S1-TH disconnecte*d) and with (*S1-TH connected*) bidirectional thalamocortical connectivity.

### 3.6 Local field potentials (LFPs) recorded from the in vivo-like S1 circuit

We simulated extracellular LFP recordings at multiple depths and horizontal distances in the S1 cortical column during the in vivo-like state (Fig. 9). The LFP calculation was based on the transmembrane currents across all segments of neurons. To reduce the computational cost of the calculation, we included only the 1,376 morphologically detailed neurons (4.4% of the total neurons) within a central cylinder of 84 μm diameter (Fig. 9A,B). The remaining 29,970 S1 and 7,266 thalamic neurons were simulated using artificial spike generators (VecStims) to ensure the dynamics of the detailed neurons were identical as in the full scale simulation (Fig. 9D). The simulated morphologically-detailed neurons therefore included the same 2,702,107 synapses with STP as those in the full in vivo simulation. We inserted recording electrodes at 4 different cortical depths (y) – 500, 1000, 1500 and 2000 μm – and 2 radial (x-z plane) distances (0 and 297 μm) from the cylinder center (Fig. 9C). Recorded LFP amplitudes were in the order 1-1000 μV consistent with the experimental literature (Michael W. Reimann et al. 2013; Hagen et al. 2018) (Figs. E,F). The amplitudes of LFPs recorded further away from the cylinder were attenuated ∼10-20x compared to those closer to the cylinder center, for example, the peak amplitudes for electrodes 1 and 5 were 401 μV and 24 μV, respectively. This is consistent with LFP amplitude being inversely proportional to the squared distance between electrode and current sources. When compared to the in vitro recorded LFPs (see Supp. Figs. S2 and S3 in Section 3 of Supplementary Material), which exhibited stronger slow frequency oscillations, the attenuation measured at the distant electrodes was only 5x. This is consistent with the observed frequency-dependent attenuation phenomenon, where high frequency signals are attenuated more than low frequency oscillations (Buzsáki, Anastassiou, and Koch 2012; Michael W. Reimann et al. 2013). The LFP power spectral densities generally depict an inverse relationship between power and frequency, which is typically described in animal LFP recordings. Overall, these preliminary results demonstrate the model can be used to simulate and capture several physiological features of extracellular LFPs.

**Figure 9.**
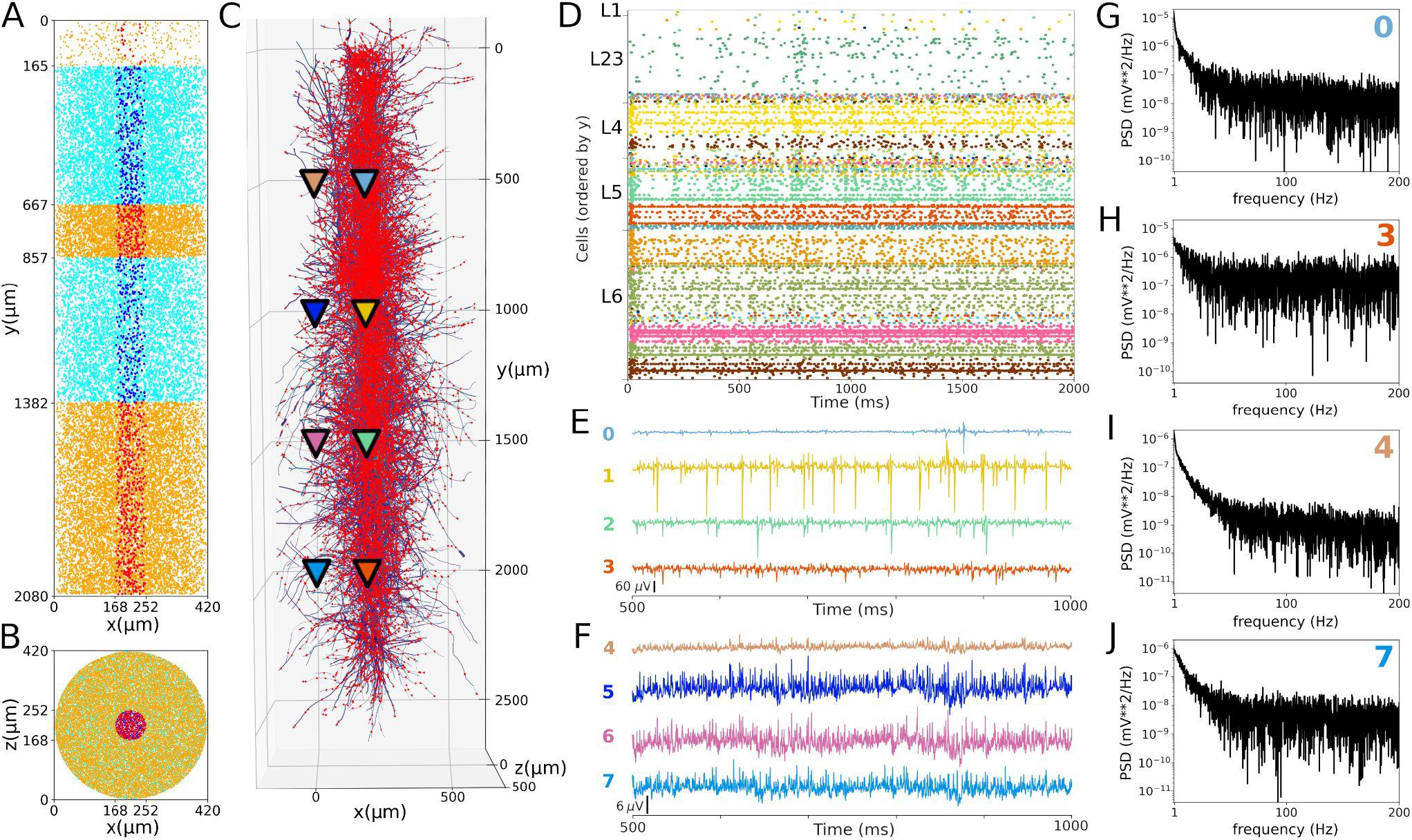
Local field potentials (LFPs) recorded from the in vivo-like S1 circuit. (A,B) Lateral and top-down 2D representation of the location of 1376 morphological cells within a cylinder with 2082 μm height and 84 μm radius. Morphologically-detailed neurons are shown in red (L1, L4, and L6) and blue (L23 and L5); whereas the 29,970 simplified cells (spike generators) are shown in orange (L1, L4, and L6) and cyan (L23 and L5) circles. (C) 3D representation of the morphologically-detailed neurons with the location of all synapses (red dots), and the location of the 8 LFP recording electrodes at 4 different depths and 2 radial distances (color triangles). (D) Spiking raster plot of the morphologically-detailed neurons used to calculate the LFP. (E) LFP signals recorded at the 4 electrodes in the center of the cylinder (colors correspond to triangles in panel C). Electrodes numbered 0, 1, 2 and 3 correspond with cortical depths (y) 500, 1000, 1500, and 2000μm, respectively. (E) Same as D, but for the LFPs recorded at a radial distance of 297μm; electrodes 4-7. (G-H) Power spectral densities (PSDs) for electrodes 0, 3, 4, and 7 calculated over a 10-second simulation (initial transient period was not included). PSDs exhibit an inverse relationship between power and frequency.

## 4 Discussion

We provided here the first large-scale S1 model that is accessible to the wider community, building on the details of the prior state-of-the-art BBP S1 model. The model closely reproduced the original cell morphologies and electrophysiological responses for the 207 morpho-electrical (me) cell types, with 5 examples for each, totaling 1035 cell models (Fig. 1); the spatial distribution of these cells across layers; and the connectivity properties of the 1941 pathways, including synaptic dynamics and short-term plasticity (Figs. 2,3). After tuning, the simulations produced reasonable dynamics with rates and activity patterns corresponding to in vivo measures of cortical activity (Figs. 5,6). There was no direct comparison to the full network dynamics of the original BBP model since original simulation data was not available. However, firing rates and overall 1 Hz underlying oscillation when using STP is comparable to that seen in the original model version paper (Markram et al. 2015) (Fig. 11). We also extended the model by adding thalamic circuits, including 6 distinct thalamic populations that reproduced cell and circuit-level dynamics, and with intrathalamic, thalamocortical and corticothalamic connectivity derived from experimental data (Fig. 4). The addition of the thalamic circuit resulted in distinct activity patterns and synchronous activity across cortical and thalamic populations (Fig. 7). Finally, we decreased the extracellular calcium concentration ([Ca^2+^]_o_) to simulate in vivo-like states with asynchronous activity (Fig. 8). Local field potentials (LFPs) recorded at multiple cortical depths and horizontal distances exhibited realistic oscillatory patterns and power spectra, including the experimentally observed distance- and frequency-dependent attenuation (Fig. 9).

The S1 model now joins other NetPyNE cortical simulations: generic cortical circuits (Romaro et al. 2021), auditory and motor thalamocortical circuits (Sivagnanam et al. 2020; Dura-Bernal, Neymotin, et al. 2022; Dura-Bernal, Griffith, et al. 2022), as well as simulations of thalamus (Moreira et al. 2021), dorsal horn of spinal cord (Sekiguchi et al. 2021), Parkinson’s disease (Ranieri et al. 2021) and schizophrenia (Metzner et al. 2020). These large cortical simulations can be extremely computer-intensive, which is a major motivation for NetPyNE’s facilities that allow one to readily simplify the network by swapping in integrate-and-fire or small-compartmental cell models, or by down-scaling to more manageable sizes. CoreNEURON is a state-of-the-art solver optimized for large scale parallel simulations, now included as part of the official NEURON package. The optimization on CPUs and the ability to run across GPUs in CoreNEURON is another key NetPyNE feature enhancing runnability. In the present case, the original S1 model is largely inaccessible, despite the cooperation of its designers, since it requires specialized tools, workflows, and training. Nonetheless, most of the data required to replicate it is available via the NMCP, which we ourselves used to implement the NetPyNE version.

We were able to get substantial speedup (>2x) for both model setup and run using CoreNEURON despite only using CPUs with no GPU at this time. We note that using CPU cycles/timestep would provide a more direct measure than the total simulation time, which may be affected by other factors such as background processes (Girardi-Schappo et al. 2017). Nonetheless, the speedup obtained is consistent with the 2-7x speedups recently reported when using CoreNEURON on CPUs to simulate large-scale models (Kumbhar et al. 2019; Awile et al. 2022). For example, the NetPyNE-based motor cortex model exhibited a speedup of 3.5x on Google Cloud. When using GPUs, speedups of up to 40x were reported. The differences in firing activity seen with NEURON vs CoreNEURON are expected due to vectorisation of the compute kernels in CoreNEURON and potential differences due to different solvers when using NMODL with sympy. Further differences are to be expected once this is extended to GPUs (Kumbhar et al. 2019; Jézéquel, Lamotte, and Saïd 2015).

We made 2 significant changes in our port to NetPyNE. First, we did not replicate the stochastic K channels that appear in 3.6% of the neurons, making our port somewhat simpler than the original. This channel required writing custom code and made simulations slower, but it could be added to the model in a future iteration. Second, we have not utilized the original cell-to-cell connection mappings that were obtained by BBP from direct microscopic observations of overlap between pre-synaptic axonal fields and post-synaptic dendritic fields (so-called Peter’s principle). In the original BBP S1 model, the use of cell-to-cell connections necessarily limited the simulation to use precisely the original model’s cell morphologies, cell positions and scales. It also required storing and loading large files of connection data. We therefore replaced this connection framework with one based on connection probability based on cell type (including layer), inter-cell distance, and dendritic pattern of post-synaptic locations. Although saving somewhat on space, there is a time-space tradeoff since this requires further calculations on start-up. Despite these limitations, we had excellent agreement with both cell model matching and connection density matching.

Our implementation also incorporates a novel model of thalamic circuitry that recapitulates multiple experimental findings at the single neuron and circuit levels. Thalamic and reticular cell models were adjusted to reproduce the reported resting membrane potential, approximately -60 and -80 mV, respectively (Jahnsen and Llinás 1984; Murray Sherman and Guillery 2009; Destexhe, Contreras, et al. 1996). In the networks, thalamocortical cells fired at low frequencies (2-4 Hz), while reticular cells fired at higher rates (6-14 Hz), consistent with values previously reported in the literature (Kim and McCormick 1998). Thalamic simulations also showed rhythmic rebound bursting when hyperpolarized and regular spiking activity at depolarized potentials (Destexhe and Contreras 2011). The thalamic network exhibited synchronous activity within and across several populations, as well as synchronous firing with cortical populations, particularly in the in vitro condition. These synchronous patterns likely emerged as a consequence of the implemented intrathalamic and thalamocortical connectivity, including the topological organization based on axonal footprints (Lam, Nelson, and Sherman 2006; Lam and Sherman 2007, 2011). Taken together, these results make the thalamic circuit a valuable extension to the S1 model, by providing a more realistic input source to the cortical circuit and enabling the study of thalamocortical interactions.

Recording the intracellular potential of multiple neurons in vivo requires an elaborate set up and is generally challenging. Extracellular recordings are more accessible and therefore more commonly used in experimental studies. Extracellular potentials are generated by transmembrane currents resulting from neuronal activity. Evidence suggests the main contributor to extracellular signals are synaptic currents (Buzsáki, Anastassiou, and Koch 2012; Michael W. Reimann et al. 2013). Computational modeling coupled with recordings of field activity in animals can provide insights into the cooperative behavior of neurons and increase our understanding of how these processes contribute to the extracellular signal (Buzsáki, Anastassiou, and Koch 2012; Michael W. Reimann et al. 2013). The simulated LFPs exhibited a similar range of amplitudes as those recorded experimentally, and reproduced several features of LFPs, including the distance-dependent and frequency-dependent attenuation. This opens the door to future validation of the model by comparing LFPs to those recorded experimentally under different conditions, and to future studies of the biophysical sources of LFPs and the exact contribution of different network populations (Hagen et al. 2018).

To place our model in the context of recent literature, we follow the classification proposed by a recent review of data-driven models structural connectivity at the microcircuit level (Shimoura et al. 2021). Our model can be classified as using conductance-based, morphologically-detailed neurons, with a network size of 38,612 neurons, synaptic plasticity and network spatiality (e.g. distance-based connectivity). Our NetPyNE implementation, together with the original BBP implementation (Markram et al. 2015), constitute the only conductance-based, morphologically-detailed models of S1. These contrast with previous models of S1 (Huang, Zeldenrust, and Celikel 2022) or of generic sensory cortex (Potjans and Diesmann 2014) that employ simpler neuron models (leaky integrate and fire point neurons). Models with detailed conductance-based and morphologically-detailed neurons have been developed for other cortical regions, including V1 (Billeh et al. 2020; Arkhipov et al. 2018), M1 (Dura-Bernal, Neymotin, et al. 2022), A1 (Dura-Bernal, Griffith, et al. 2022), and CA1 (Bezaire et al. 2016; Ecker et al. 2020). Our model is also unique in incorporating thalamic neurons and thalamocortical bidirectional topological connectivity. Previous thalamocortical circuit models included less biophysically-detailed neuron models and simpler connectivity (Izhikevich and Edelman 2008), or focused on single cell (Iavarone et al. 2019) or small circuit models (Murray and Anticevic 2017). An impressively detailed model of the thalamoreticular microcircuit has recently been developed, although this is limited to the VPL and RTN somatosensory thalamus regions (Iavarone et al. 2022).

As outlined above, the level of biophysical, morphological and connectivity detail in the model is very high compared to most existing models. Although this makes it harder to simulate and tune, it also enables exploration of a unique set of scientific questions that simpler models cannot address, or at least not with the same level of realism. Here we included two results that require and justify the level of detail of the model. First, we simulated a network state with lower extracellular calcium concentration that more closely resembles the in vivo conditions (Fig. 8). Secondly, we calculated realistic LFPs, which critically depend on the sum of transmembrane currents along detailed neuronal morphologies (Fig. 9). We also describe the methodology for future model parameter explorations, and provide a basic code set up example to explore the effects of inhibitory GABAergic connections on network dynamics. Examples of parameter explorations in NetPyNE-based biophysically detailed circuit models can be found in our related publications on motor and auditory cortex models (Sivagnanam et al. 2020; Dura-Bernal, Neymotin, et al. 2022; Dura-Bernal, Griffith, et al. 2022), including an exploration of the effects of long-range and neuromodulatory inputs.

Consequently, our port of the S1 model provides a quantitative framework that can be used in several ways. First, it can be used to perform in silico experiments to explore sensory processing under the assumption of various coding paradigms or brain disease, including the representation of whisker motion (Bosman et al. 2011; Huang, Zeldenrust, and Celikel 2022), maximization of sensory dynamic range (Gautam et al. 2015), response to unexpected sensory inputs, schizophrenia (Metzner et al. 2020) and Parkinson’s disease (Ranieri et al. 2021). Second, drug effects can be directly tested in the simulation (Neymotin et al. 2016) -- this is an advantage of a multiscale model with scales from molecule to network, which is not available in simpler models that elide these details. Third, the model constitutes a unified multiscale framework for organizing our knowledge of S1 which serves as a dynamical database to which new physiological, transcriptomic, proteomic, and anatomical data can be added. This framework can then be utilized as a community tool for researchers in the field to test hypotheses and guide the design of new experiments.

## Acknowledgments

This work was funded by the following grants: NIH NIBIB U24EB028998, NSF 1904444-1042C, NYS SCIRB DOH01-C32250GG-3450000 and NIH NIDCD R01DC012947. This research was funded in part by Aligning Science Across Parkinson’s [ASAP-020572] through the Michael J. Fox Foundation for Parkinson’s Research (MJFF). For the purpose of open access, the author has applied a CC BY public copyright license to all Author Accepted Manuscripts arising from this submission.

## 1 Data Availability Statement

The model and data for this study can be found in the following repositories and online platforms: GitHub (github.com/suny-downstate-medical-center/S1_netpyne/), ModelDB (link), Open Source Brain (link) and EBRAINS Model Catalog (link).

