## Supplementary Material for "Large-scale biophysically detailed model of somatosensory thalamocortical circuits in NetPyNE"

### 1 Comparison of cells with StochKv channels in NetPyNE and BBP implementations

The NetPyNE implementation employs a deterministic version of the stochastic potassium channel (StochKv) channel. Neurons with the deterministic StochKv channel exhibited lower spiking irregularity (Fig. S1). This occurred primarily when the neurons received a weak stimulus (Figs. S1A,C,D), since in that case the StochKv channel has a stronger effect. However, for moderate-strong stimulation, the neuronal responses using the stochastic vs the deterministic version of the channel are very similar (Figs. S1B,E,F). Furthermore, in the context of the full network simulation, cells with the deterministic version of StochKv exhibit irregular spiking patterns (Fig. S1G).

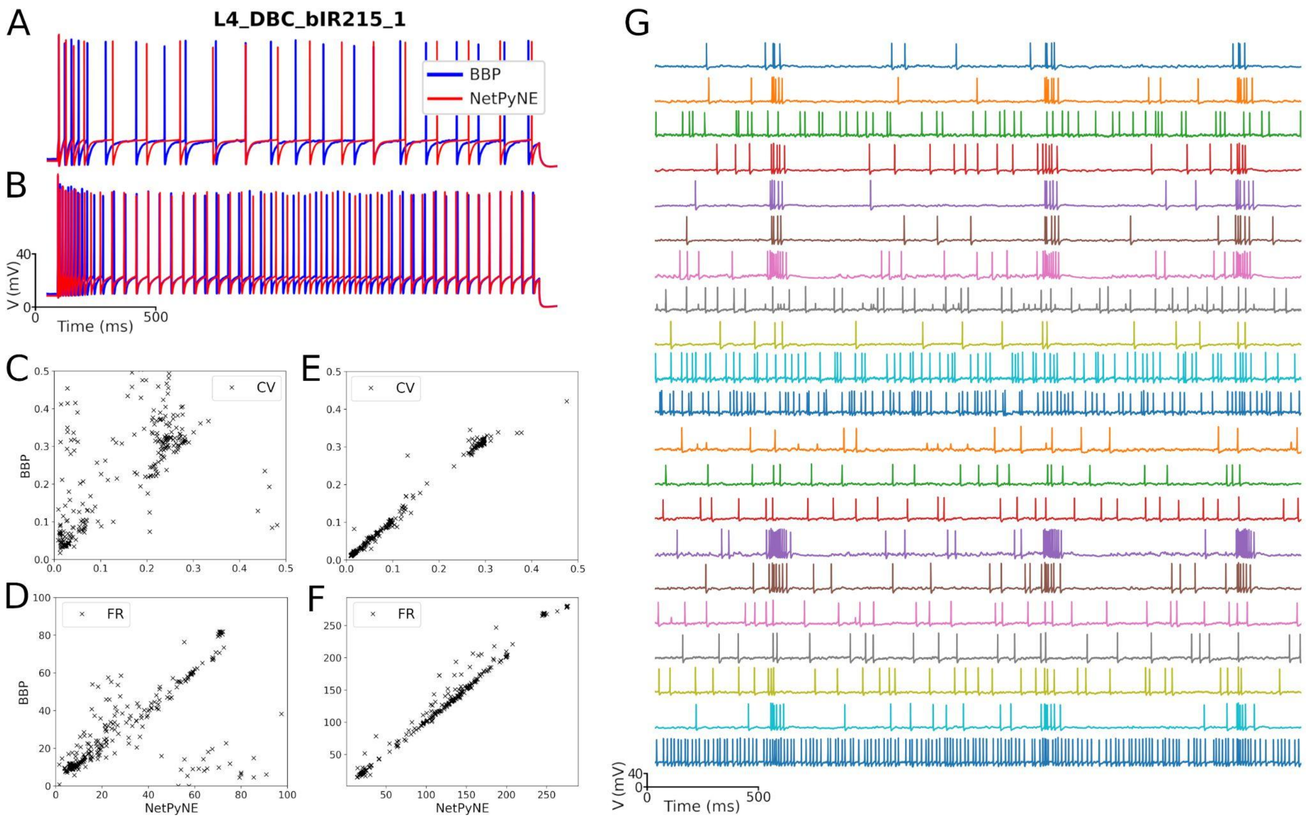


**Figure S1. Cells with StochKv channels in NetPyNE vs BBP.** (A,B) Somatic membrane potential of the neurons under current clamp with amplitude 0.1 nA and 0.8 nA, respectively. NetPyNE results (red) compared to the original BBP model results (blue) for the L4_DBC_bIR215_1 cell. The deterministic version of the BBP stochastic potassium channel (StochKv) best matches the traces in the high current case. (C,D) Comparison of BBP's and NetPyNE's inter spike interval coefficient of variation (CV) and firing rate (FR) with amplitude 0.1 nA during 2 seconds for each cell type. (E,F) Same comparison as in C,D but using 0.8 nA. (G) Example traces of cells with the StochKv in the NetPyNE full circuit simulations (Figure 6).

### 2 Parameter optimization/exploration

Although most model parameters are constrained by experimental data, it is typically necessary to perform parameter optimization (also referred to as tuning) to ensure the model reproduces experimental measures such as population firing rates. Once the model has been optimized and validated against experimental data, researchers can explore model parameters to ask questions about the modeled system, gain insights and generate hypotheses and predictions. Parameter optimization and exploration is typically done by systematically modifying the value of certain parameters (e.g. ionic channel conductances or synaptic weights). As previously described, NetPyNE automates the optimization and exploration of parameters. The user can easily define the range of values to explore and the simulations set up using a simple format based on Python dictionaries and lists, or the graphical interface. NetPyNE includes built-in customizable set ups for different environments, multicore machines with MPI, HPCs with SLURM or HPC/Torque workload managers, etc. Once the user runs the parameter exploration/exploration, NetPyNE will automatically submit all the required simulation jobs for each of the parameter combinations. The tool includes several other features to facilitate simulation management, including automatic filename generation, optional user-defined labels to track model versions and batch simulations, and standardized structure for output files that includes model version, NetPyNE version, parameters, network instance and simulation output data (Dura-Bernal et al. 2019).

The parameters optimized for the S1 thalamocortical circuit were the background rate for excitatory and inhibitory connections. For the intrathalamic projections, we optimized connection weight (range 0 to 2 mV), connection probability (range 0 to 1), y-axis connection radius (1, 2, 5 or 10%) and connectivity divergence of the HO populations (5, 10, 20 or 40 cells). For the thalamocortical and corticothalamic projections we optimized connection weight (range 0 to 2 mV) and connection probability (range 0 to 1).

Although performing a detailed model parameter exploration is out of the scope of this paper, here we provide an example set up and describe the basic steps to follow. In the example we aim to explore the effect of inhibitory GABAergic synaptic strength on network dynamics. We therefore selected the two relevant parameters: IEGain and IIGain, the gain factors for the synaptic strength of I→E and I→I connections. These parameters were already defined in the cfg.py file, with a default value of 1.0, and are used in netParams.py to scale the weights of the relevant connections. A reasonable range of values to explore initially might be [0.5, 0.75, 1.0, 1.25, 1.5] for each parameter – these can be adjusted iteratively based on simulation results. In order to quantify the robustness and validate our results statistically, we will simulate each parameter combination using 5 connectivity and 5 input randomization seeds. The code to set up the described parameter exploration is available here: https://github.com/suny-downstate-medical-center/S1_netpyne/blob/master/sim/batch.py (inhib() function). After executing the parameter exploration, NetPyNE will run 625 simulations corresponding to the 5*5*5*5 (IEGain, IIGain, connectivity seed, input seed) parameter value combinations. Researchers can then analyze and visualize any of the network output measures, from voltage traces to mean firing rates to LFP power, to study the effect of inhibitory connection strength on network activity.

### 3 Local Field Potential recorded from the in vitro-like S1 circuit

The in vitro-like simulation with bidirectional thalamic connectivity (Fig. 7) exhibits activity with bursts synchronization. In the same way as the in vivo-like case (Fig. 9), the microcircuit has 1376 morphological, 29,970 S1 spike cells, 7,266 thalamic spike cells, 499,412 connections, and 2,702,107 synapses modeled with STP. LFP recording electrodes were located in the cylinder center (x=z=210µm) and at a radial distance from the center (x=z=0µm), at 3 different depths (1000, 1200, and 1400µm). A representation of the network cells, recording electrodes, and the 5-second spiking raster plot is shown in Figure S2. The corresponding LFP recorded signals are shown in Figure S3, which also illustrates the observed distance- and frequency-dependent attenuation (Fig. S3A-D) and the inverse relation of power and frequency (Fig. S3E-F).

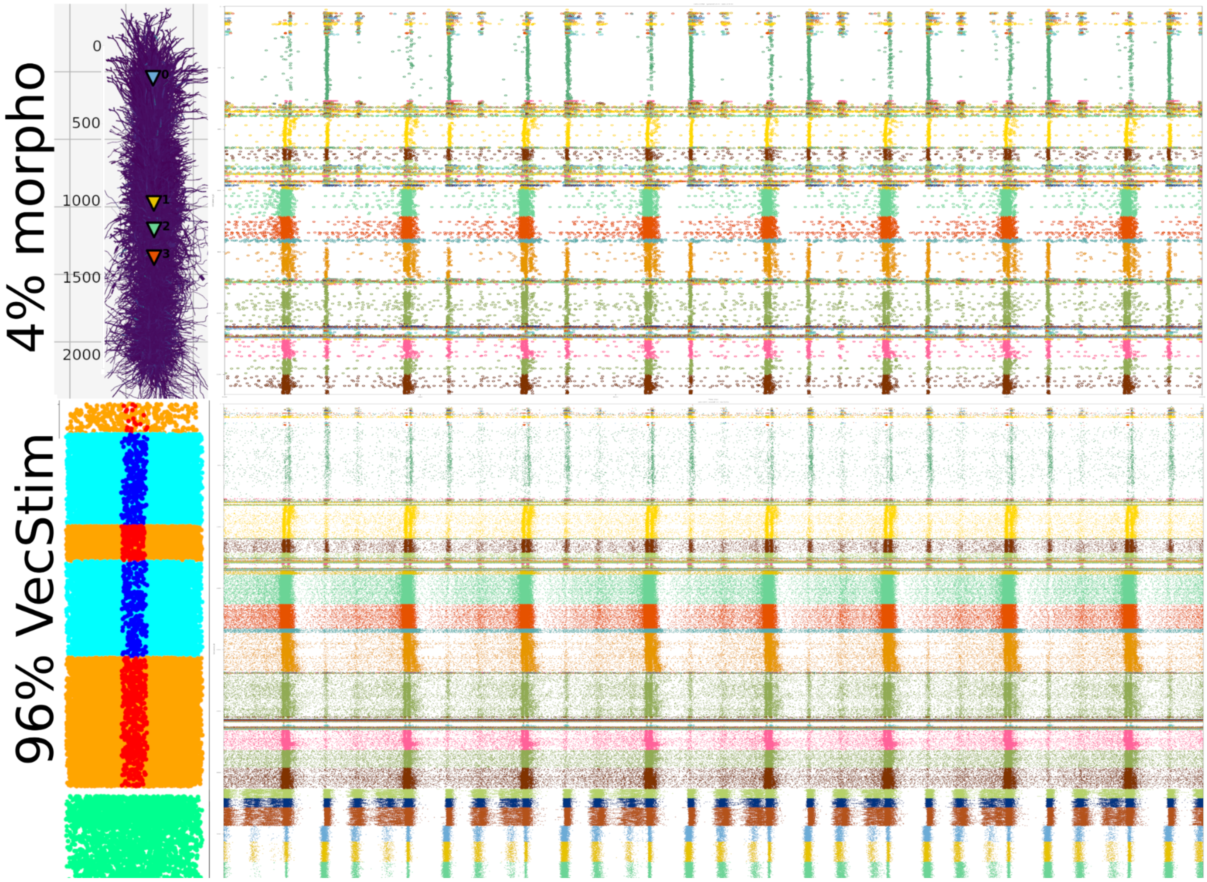


**Figure S2. Subsampled S1 model used to calculate LFPs.** Spiking raster plots of the S1 model related to Fig. 7. Activity from the 4.4% of neurons simulated using detailed morphological reconstructions (top panel) and the 95.6% of inputs simulated using spike generators with spike times recorded from a previous simulation.

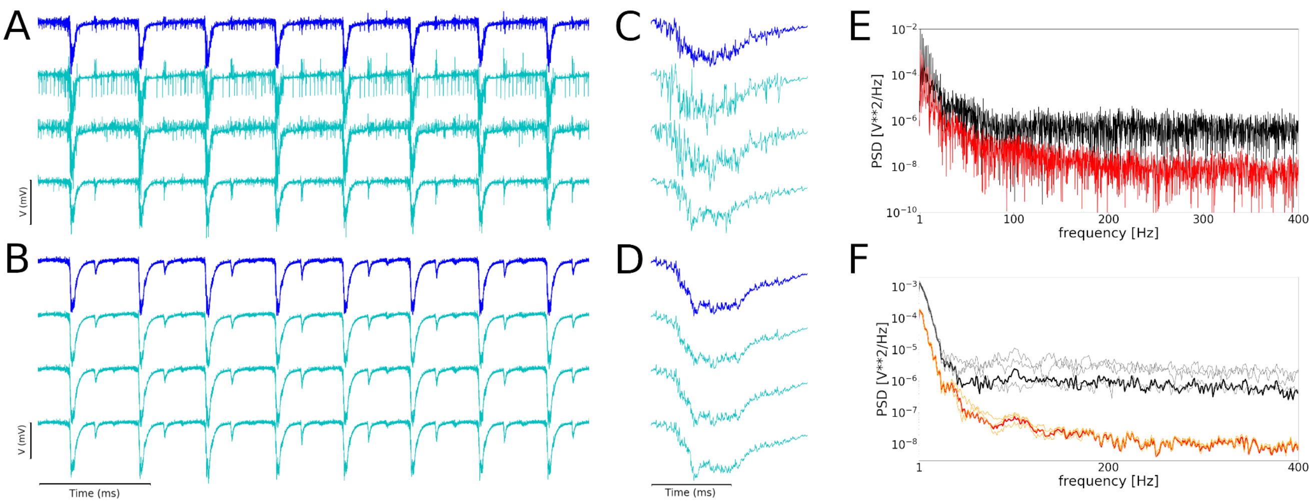


**Figure S3. Local field potentials (LFPs) recorded from the in vitro-like S1 circuit.** S1 model LFP signal for the 4.4% neurons simulated using detailed morphological reconstructions recorded from (A) electrodes located in center of cylinder (bar 0.5 mV), and (B) electrodes located at a radial distance (x=z=210 µm) from cylinder center (bar 0.1 mV). The horizontal bar represents 1000 ms. (C,D) Zoomed in representation of LFP signal, withe horizontal bar representing 50 ms. The blue traces represent the mean LFP signal. (E) Periodogram for LFP recorded from deepest electrode at the central (black) and radially distant (red) locations; corresponding to bottom plots in A and B, respectively. (F) Welch frequency plot for all traces in A (black and gray lines) and B (red and orange lines).

### 4 Acronyms

The full name of cells' m-types, s-types and e-types and their corresponding acronyms are summarized in Table S1. The full name and acronym of the 207 cell types can be found in the Neocortical Microcircuit Collaboration (NMCP; https://bbp.epfl.ch/nmc-portal).

| m-types | |
|---|---|
| **DAC** | Descending Axon Cell |
| **NGC-DA** | Neurogliaform Cell with dense axonal arborization |
| **NGC-SA** | Neurogliaform Cell with slender axonal arborization |
| **HAC** | Horizontal Axon Cell |
| **LAC** | Large Axon Cell |
| **SAC** | Small Axon Cell |
| **MC** | Martinotti Cell |
| **BTC** | Bitufted Cell |
| **DBC** | Double Bouquet Cell |
| **BP** | Bipolar Cell |
| **NGC** | Neurogliaform Cell |
| **LBC** | Large Basket Cell |

| | |
|---|---|
| **NBC** | Nest Basket Cell |
| **SBC** | Small Basket Cell |
| **ChC** | Chandelier Cell |
| **PC** | Pyramidal Cell |
| **SP** | Star Pyramidal Cell |
| **SS** | Spiny Stellate Cell |
| **TTPC1** | Thick-tufted Pyramidal Cell with a late bifurcating apical tuft |
| **TTPC2** | Thick-tufted Pyramidal Cell with an early bifurcating apical tuft |
| **UTPC** | Untufted Pyramidal Cell |
| **STPC** | Slender-tufted Pyramidal Cell |
| **TPC** | Tufted Pyramidal Cell |
| **IPC** | Pyramidal Cell with inverted apical-like dendrites |
| **BPC** | Pyramidal Cell with bipolar apical-like dendrites |
| **s-types** | |
| **I1** | Inhibitory facilitating |
| **I2** | Inhibitory depressing |
| **I3** | Inhibitory pseudo-linear |
| **E1** | Excitatory facilitating |
| **E2** | Excitatory depressing |
| **E3** | Excitatory pseudo-linear |
| **e-types** | |
| **cADpyr** | continuous Accommodating (Adapting) for pyramidal cells |
| **cAC** | continuous Accommodating |
| **bAC** | burst Accommodating |
| **cNAC** | continuous Non-accomodating |
| **bNAC** | burst Non-accomodating |
| **dNAC** | delayed Non-accomodating |
| **cSTUT** | continuous Stuttering |
| **bSTUT** | burst Stuttering |
| **dSTUT** | delayed Stuttering |
| **cIR** | continuous Irregular |
| **bIR** | burst Irregular |

**Table S1. Glossary.** m-type: morphological type; s-type synapse type; e-type: electrical type. Note that the cell type L1_DLAC refers to the LAC m-type of layer 1.
